# Phylogenomic analyses of the East Asian endemic *Abelia* (Caprifoliaceae) shed insights into the temporal and spatial diversification history with widespread hybridizations

**DOI:** 10.1101/2021.04.13.439739

**Authors:** Qing-Hui Sun, Diego F. Morales-Briones, Hong-Xin Wang, Jacob B. Landis, Jun Wen, Hua-Feng Wang

**Affiliations:** Key Laboratory of Tropical Biological Resources of Ministry of Education, College of Tropical Crops, Hainan University, Haikou 570228, China; Department of Plant and Microbial Biology, College of Biological Sciences, University of Minnesota, 140 Gortner Laboratory, 1479 Gortner Avenue, Saint Paul, MN 55108, USA; Systematics, Biodiversity and Evolution of Plants, Department of Biology I, Ludwig-Maximilians-Universität München, Menzinger Str. 67, 80638, Munich, Germany; Zhai Mingguo Academician Work Station, Sanya University, Sanya 572022, China; School of Integrative Plant Science, Section of Plant Biology and the L.H. Bailey Hortorium, Cornell University, Ithaca, NY 14850, USA; BTI Computational Biology Center, Boyce Thompson Institute, Ithaca, NY 14853, USA; Department of Botany, National Museum of Natural History, MRC-166, Smithsonian Institution, PO Box 37012, Washington, DC 20013-7012, USA

**Keywords:** *Abelia*, Gene tree discordance, Hybridization, Mainland China-Taiwan island disjunction

## Abstract

**Background and Aims:** *Abelia* (Caprifoliaceae) is a small genus with five species, including one man-made hybrid and several natural hybrids. The genus has a discontinuous distribution in mainland China, Taiwan Island, and the Ryukyu islands, providing a model system to explore mechanisms of species dispersal in the East Asian flora. However, the current phylogenetic relationships within *Abelia* remain uncertain.

**Methods:** In this study, we reconstructed phylogenetic relationships within *Abelia* using nuclear loci generated by target enrichment and plastomes from genome skimming. Divergence time estimation, ancestral area reconstruction, and ecological niche modelling (ENM) were used to examine the diversification history of *Abelia*.

**Key Results:** We found extensive cytonuclear discordance across the genus. By integrating lines of evidence from molecular phylogenies, divergence times, and morphology, we propose to merge *A. macrotera* var. *zabelioides* into *A. uniflora.* Network analyses suggested that there have been widespread and multiple hybridization events among *Abelia* species. These hybridization events may have contributed to the speciation mechanism and resulted in a high observed morphological diversity. The diversification of *Abelia* began in the early Eocene, followed by *A. chinensis* var. *ionandra* colonizing the island of Taiwan in the Middle Miocene. The ENM results suggested an expansion of climatically suitable areas during the Last Glacial Maximum and range contraction during the Last Interglacial. Disjunction between the Himalayan-Hengduan Mountain region (HHM) and the island of Taiwan is most likely the consequence of topographic isolation and postglacial contraction.

**Conclusions:** We used genomic data to reconstruct the phylogeny of *Abelia* and found a clear pattern of reticulate evolution in the group. In addition, our results support shrinkage of postglacial range and the heterogeneity of the terrain have led to the disjunction of the mainland China-Taiwan island. This study provides important new insights into the speciation process and taxonomy of *Abelia*.

## INTRODUCTION

The underlying causes of disjunct distributions in various organisms have been a central issue in biogeography (e.g., Xiang et al., 1998; Wen, 2001; Qian and Ricklefs, 2004; Wang et al., 2012; Wen et al., 2013, 2018). Much discussion has focused on two mechanisms, vicariance and long-distance dispersal (Givnish and Renner, 2004; Yoder and Nowak, 2006; Šarhanová et al., 2017; Wang et al., 2020b). The emergence of geographic barriers leads to vicariance, resulting in decreased gene flow; while long-distance dispersal in plants usually involves rare events driven by dispersal vectors such as animals, wind or water currents (Nathan et al., 2008). Recent molecular systematic studies have shown that many plant disjunctions are too young to be explained by continental drift or major geographic barriers (Mummenhoff et al., 2004; Yoder and Nowak, 2006; Linder et al., 2013; Jiang et al., 2019). Hence, long-distance dispersal has emerged as a major mechanism to explain many discontinuous distributions in historical biogeography (Wen et al., 2013; Ye et al., 2019; Wen and Wagner, 2020).

Biogeographic disjunctions between mainland China-Taiwan island have received considerable attention with the separation of the two regions by 130 km (Wu and Wang, 2001; Xiang and Soltis, 2001; Qiu et al., 2011; Chen et al., 2012; Ye et al., 2014, Jin et al., 2020). The island of Taiwan is the largest subtropical mountainous island in the monsoonal western Pacific region, located on the Tropic of Cancer, having originated in the late Miocene (ca. 9-5 million years ago; Sibuet and Hsu, 2004). Molecular phylogenetic analyses have shown that several plant lineages have disjunct distributions between mainland China and Taiwan Island (e.g., *Sassafras* J. Presl, Nie et al., 2007; *Pseudotsuga* Carrière, Wei et al., 2010; *Triplostegia* Wall. ex DC., Niu et al., 2018; *Prinsepia*., Jin et al., 2020). Geologically, the island of Taiwan and its adjacent continental margin of mainland China belong to the Eurasian plate; in the Late Cretaceous both were part of the continental block. The continental block has undergone many extensions and rifting events since the Paleocene (Wei, 2010; Suo et al., 2015). Two hypotheses have been proposed to explain the formation of current plant disjunctions in this region: long-distance dispersal and postglacial contraction (Chou et al., 2011; Wang et al., 2013; Ye et al., 2014; Niu et al., 2018). A phylogeographic study of *Cunninghamia* R.Br. ex A.Rich. (Taxodiaceae) revealed that populations of Taiwan derived from continental Asia and colonized the island via long-distance seed dispersal (Lu et al., 2001). Factors that have major impacts on population dynamics and genetic variation patterns in plants in this region include past geological and climatic changes, uplift of the Qinghai-Tibet Plateau, global optimum temperature since the Miocene and the Quaternary glacial-interglacial transition (Xu et al., 2015; Cao et al., 2016; Chen et al., 2020). Recent studies have suggested that although the ice sheet was not ubiquitous from the Qinling Mountains and Huai River (at ca. 34° N) to the tropical south (≤22° N) in China during the Last Glacial period, the temperature in most parts of mainland China was lower than today (Shi et al., 2006). Temperature changes influence species distributions, population sizes, and intraspecific genetic differentiation. The postglacial contraction hypothesis states that many species had a continuous distribution from mainland China to Taiwan prior to the Quaternary glacial period, sometimes with disjunction between the Himalayan-Hengduan Mountain region and the island of Taiwan occurring when samples migrated to higher altitudes as global temperatures increased.

*Abelia* R.Br. belongs to the subfamily Linnaeoideae Raf. within Caprifoliaceae *s.l.* (Dipsacales; Yang and Landrein, 2011; Caprifoliaceae; Wang et al., 2020a, 2021). The genus includes *A. chinensis* (four varieties), *A. forrestii* (two varieties), *A.* × *grandiflora*, *A. macrotera* (eight varieties), *A. schumannii* and *A. uniflora* (Landrein et al., 2017; 2020; Fig. 1). *Abelia* is a typical East Asian endemic genus with a disjunct distribution in southwestern China, extending to the eastern portion of the island of Taiwan and the Ryukyu islands in Japan. The genus includes an important ornamental species, *Abelia* × *grandiflora*, widely cultivated in yards, parks and roadsides of temperate cities (e.g., Beijing, London, Washington D.C., and Wuhan). However, the phylogenetic relationships within *Abelia* remain unresolved due to limited DNA markers and sampling, and hence the biogeographic history and process of speciation are still unclear. Kim (1998) used one nuclear (ITS) and one plastid (*matK*) marker to infer the phylogenetic relationships within *Abelia*. However, species relationships could not be clarified because only two *Abelia* species were included, and a polytomy existed between the two *Abelia* species and those of *Diabelia*. Three *Abelia* species were included (*A. chinensis*, *A. engleriana* and *A. schumannii*) in Jacobs et al. (2010), but no information about relationships can be inferred since the three formed a polytomy. Using ITS and nine plastid regions, Wang et al. (2015) found that *A. forrestii* was sister to all other *Abelia* species, while *A. chinensis* was sister to *A.* × *grandiflora* with moderate support. Although Landrein (2017) included 46 samples (representing five species) of *Abelia* (22 from natural samples and 24 of horticultural breeds), using AFLP, ITS and three plastid genes, the species phylogeny was still unclear due to lack of resolution and low bootstrap support for many nodes.

**Figure 1.**
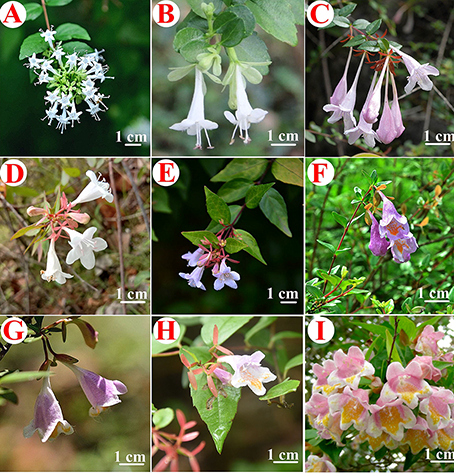
Photographs of *Abelia* taxa. (A) *A. chinensis*; (B) *A. chinensis* var. *ionandra*; (C) *A. forrestii* (photograph by Sven Landrein); (D) *A*. × *grandiflora*; (E) *A*. × *grandiflora* ‘Francis Mason’ (photograph by Sven Landrein); (F) *A. macrotera*; (G) *A. macrotera* var. *macrotera* (photograph by Sven Landrein); (H) *A. uniflora* (photograph by Pan Li); (I) *A. schumannii* (photograph by Sven Landrein).

Here, we explore the phylogenetic relationships and biogeographic diversification of *Abelia* using a target enrichment data set for nuclear loci and plastomes from genome skimming. We aim to (1) clarify the species-level phylogeny of *Abelia* based on nuclear genes and plastome data; and (2) reconstruct the biogeographic history of *Abelia* and explore the evolution of its distribution patterns related to the disjunction between mainland China and the island of Taiwan.

## MATERIALS AND METHODS

### Sampling, genomic data generation

We sampled 38 individuals from four species and one hybrid of *Abelia* (Landrein, 2017; 2020). Six Linnaeoideae species [*Diabelia serrata* (Siebold & Zucc.) Landrein, *D. sanguinea* (Makino) Landrein, *Kolkwitzia amabilis* Graebn., *Dipelta floribunda* Maxim. (two individuals), *Linnaea borealis* L., and *Vesalea occidentails* Villarreal] were used as outgroups. Vouchers of newly collected samples were deposited in the herbarium of the Institute of Tropical Agriculture and Forestry (HUTB), Hainan University, Haikou, China and Kew (K). Complete voucher information is listed in Supporting Information Table S1.

DNA was extracted from dried leaf tissue using a modified cetyltrimethylammonium bromide (CTAB) method, with chloroform: isoamyl alcohol separation and isopropanol precipitation at −20 Doyle and Doyle, 1987). We checked the concentration of each extraction with a Qubit 2.0 Fluorometer (Thermo Fisher Scientific, Waltham, MA, USA) and sonicated 400 ng of DNA using a Covaris S2 (Covaris, Woburn, MA) to produce fragments ∼150 - 350 bp in length for making sequencing libraries. To ensure that genomic DNA was sheared to the appropriate fragment size, we evaluated all samples on a 1.2% (w/v) agarose gel.

We used a Hyb-Seq approach (Weitemier et al., 2014) combining target enrichment and genome skimming, which allows simultaneous data collection for low-copy nuclear genes and high-copy genomic targets. To obtain nuclear data from target enrichment, we ran MarkerMiner v.1.2 (Chamala et al., 2015) with default settings to identify putative single copy nuclear (SCN) genes using the transcriptome of *Lonicera japonica* from 1KP (He et al., 2017), and the genome of *Arabidopsis thaliana* (L.) Heynh. (Gan et al., 2011). We then used GoldFinder (Vargas et al., 2019) to further filter the SCN genes identified by MarkerMiner. Finally, we obtained 428 putatively single-copy orthologous genes in total for phylogenetic analysis (Wang et al., 2021). Hybridization, enrichment, and sequencing methods followed those of Wang et al. (2021). We obtained full plastomes from genome skimming data (Wang et al., 2021) with detailed descriptions of plastome sequencing methods in Wang et al. (2020a, 2020b).

### Read processing and assembly

Raw sequencing reads were cleaned using Trimmomatic v.0.36 (Bolger et al., 2014) by removing adaptor sequences and low-quality bases (ILLUMINACLIP: TruSeq_ADAPTER: 2:30:10 SLIDINGWINDOW: 4:5 LEADING: 5 TRAILING: 5 MINLEN: 25). Nuclear loci were assembled with HybPiper v.1.3.1 (Johnson et al., 2016). Individual exon assemblies were used to avoid chimeric sequences in multi-exon genes produced by potential paralogy (Morales-Briones et al., 2018) and only exons with a reference length of more than 150 bp were assembled. Paralog detection was performed for all exons with the ‘paralog_ investigator’ warning. To obtain ‘monophyletic outgroup’(MO) orthologs (Yang and Smith, 2014), all assembled loci (with and without paralogs detected) were processed following Morales-Briones et al. (2021).

For plastome assembly, raw reads were trimmed using SOAPfilter_v2.2 (BGI, Shenzhen, China). Adapter sequences and low-quality reads with Q-value ≤20 were removed. Clean reads were assembled against the plastome of *Kolkwitzia amabilis* (KT966716.1) using MITObim v.1.8 (Hahn et al., 2013). Assembled plastomes were preliminarily annotated with Geneious R11.04 (Biomatters Ltd., Auckland, New Zealand) to detect possible inversions/rearrangements, with corrections of start/stop codons based on published *Abelia* plastomes (*A. chinensis*, accession No. MN384463.1; *A. forrestii*, accession No. MN524636.1; *A. × grandiflora*, accession No. MN524635.1; *A. macrotera*, accession No. MN524637.1).

### Phylogenetic analyses

#### Nuclear data set

We analyzed the nuclear data set using concatenation and coalescent-based methods. Individual exons were aligned with MAFFT v.7.037b (Katoh and Standley, 2013) and aligned columns with more than 90% missing data were removed using Phyutility (Smith and Dunn, 2008) prior to concatenation. Maximum likelihood (ML) analysis was conducted using IQ-TREE v.1.6.1 (Nguyen et al., 2015) while searching for the best partition scheme (Lanfear et al., 2012) followed by ML gene tree inference and 1000 ultrafast bootstrap replicates (Hoang and Chernomor 2018). Additionally, we used Quartet Sampling (QS; Pease et al., 2018) to distinguish strong conflict from weakly supported branches. We carried out QS with 1000 replicates. We used ASTRAL-III v.5.7.1 (Zhang et al., 2018) for species tree estimation, with the input being individual exon trees generated with RAxML using a GTRGAMMA model.

#### Plastome data set

Contiguous contigs were aligned with MAFFT v.7.407 (Katoh and Standley, 2013) and columns with more than 90% missing data were removed using Phyutility. We performed extended model selection (Kalyaanamoorthy et al., 2017) followed by ML gene tree inference and 200 non-parametric bootstrap replicates for branch support in IQ-TREE. Additionally, we evaluated branch support with QS using 1,000 replicates.

### Species network analysis

Due to computational restrictions and given our primary focus on potential reticulation between species, we reduced the 45 taxa data set to one outgroup and 14 ingroup samples representing all major clades based on the nuclear analyses. We looked for evidence of hybridization within *Abelia* via species network searches carried out with the InferNetwork_MPL function in PhyloNet (Wen et al., 2018). Network searches were carried out using only nodes in the gene trees that had at least 50 percent bootstrap support, enabling up to five hybridization events and optimizing the branch lengths and probabilities of inheritance of the returned species networks under the full probability. To disentangle nested hybridization, we also created a reduced, 12 sample data set by removing three known hybrids (*A.*× *grandiflora* E159, *A.*× *grandiflora* E227 and *A. chinensis* E206; see results) and carried out analyses as previously described. We used the command ‘CalGTProb’ in PhyloNet to calculate the probability scores of the concatenated tree, ASTRAL, and plastid trees to estimate the optimal number of hybridization events and to test if the network model explained our gene trees better than a purely bifurcating tree. We selected the best-fit-species network model using the lowest values for bias-corrected Akaike information criterion (AICc; Sugiura, 1978), Akaike information criterion (AIC; Akaike, 1998), and the Bayesian information criterion (BIC; Schwarz, 1978).

### Divergence time estimation

To examine the historical biogeography of *Abelia*, we performed a dated phylogenetic analysis using BEAST v.1.8 (Drummond et al., 2012). As there is potential ancient hybridization in *Abelia*, we estimated dates separately using the nuclear and plastid data. We selected the fruit fossil of *Diplodipelta* from the late Eocene Florissant flora of Colorado (Manchester, 2000; Bell and Donoghue, 2005) to calibrate the node containing *Diabelia* and *Dipelta*, as the fossil shows morphological similarities to those two genera. We set a lognormal prior for the divergence at the stem of *Diabelia* and its sister clade *Dipelta* (offset 36 Ma, log-normal prior distribution 34.07-37.20 Ma). The ancestral node of Linnaeoideae (the root of our tree) was constrained to 50.18 Ma (mean 50.18 Ma) from results of previous studies (Wang et al., 2021) despite the lack of fossil evidence. The BEAST analyses were performed using an uncorrelated log normal (UCLN) relaxed clock with a Yule tree prior, a random starting tree, and GTR + G as model of sequence evolution (as selected by MrModelTest v.2.3; Nylander et al., 2004). Each MCMC run was conducted for 100,000,000 generations with sampling every 5,000 generation. The first 10% of trees were discarded as burn-in. We used Tracer v.1.6 to assess convergence and effective sample size (ESS) > 200 (Rambaut et al., 2014). Tree Annotator v.1.8.0 (Drummond et al., 2012) was used to summarize the set of post-burn-in trees and their parameters to produce a maximum clade credibility chronogram showing the mean divergence time estimates with 95% highest posterior density (HPD) intervals.

### Ancestral area reconstructions

We reconstructed the historical biogeography of *Abelia* using Bayesian binary MCMC (BBM) analysis implemented in RASP v.3.21 (Yu et al., 2015). To account for phylogenetic uncertainty, 5,000 out of 20,000 post-burn-in trees from the BEAST analyses (nuclear data) were randomly chosen for the BBM analyses. We compiled distribution data for the species of *Abelia* and assigned the included taxa to their respective ranges. Considering the areas of endemism in *Abelia* and tectonic history of the continent, we used three areas: (A) southwestern China, (B) central-eastern China, and (C) Taiwan and Ryukyu islands. Each sample was assigned to its respective area according to its contemporary distribution range (Table 1). We set the root distribution to null, applied 10 MCMC chains each run for 100 million generations, sampling every 10,000 generations using the JC + G model. The MCMC samples were examined in Tracer v.1.6 (Drummond et al., 2012) to confirm sampling adequacy and convergence of the chains to a stationary distribution.

**Table 1.**
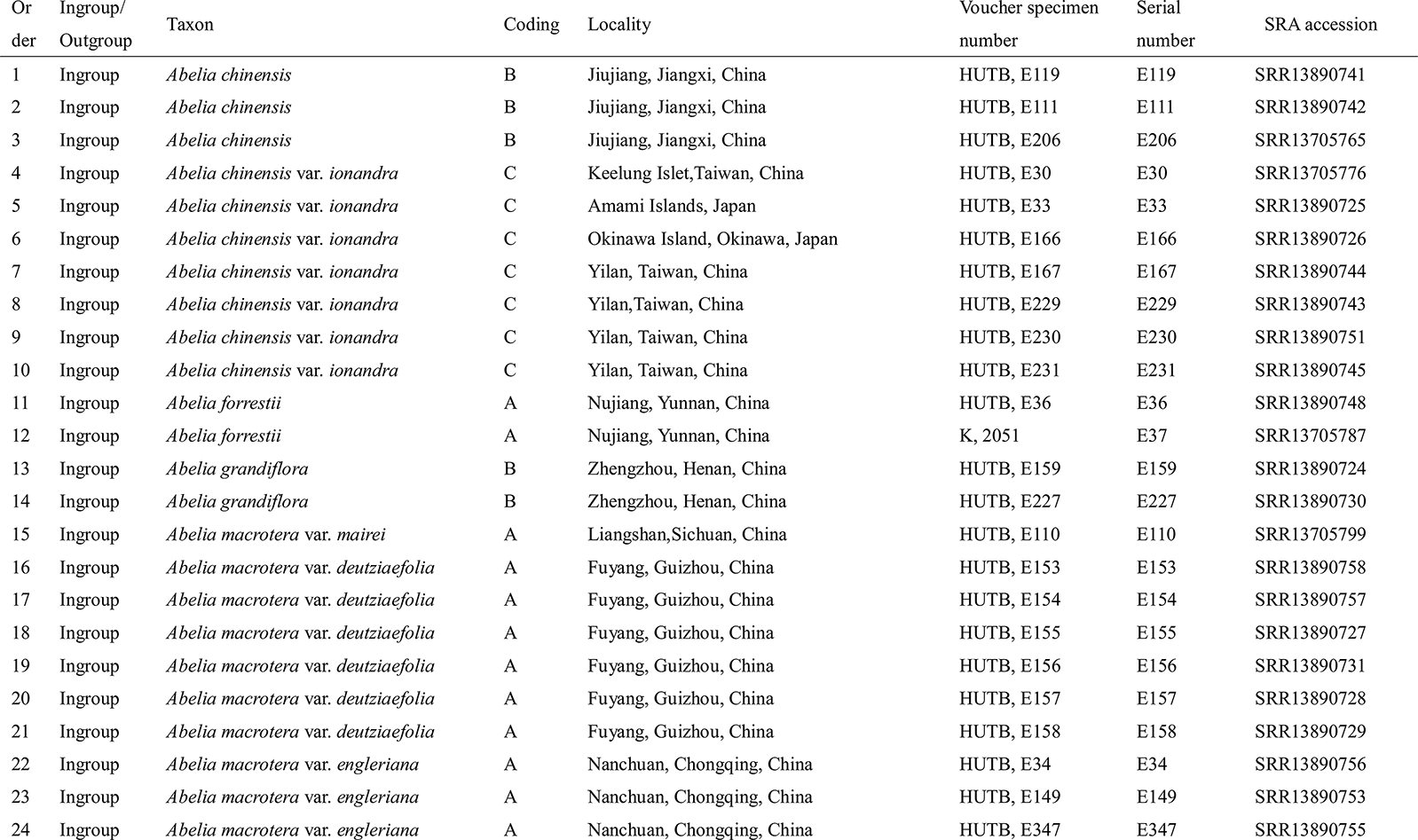
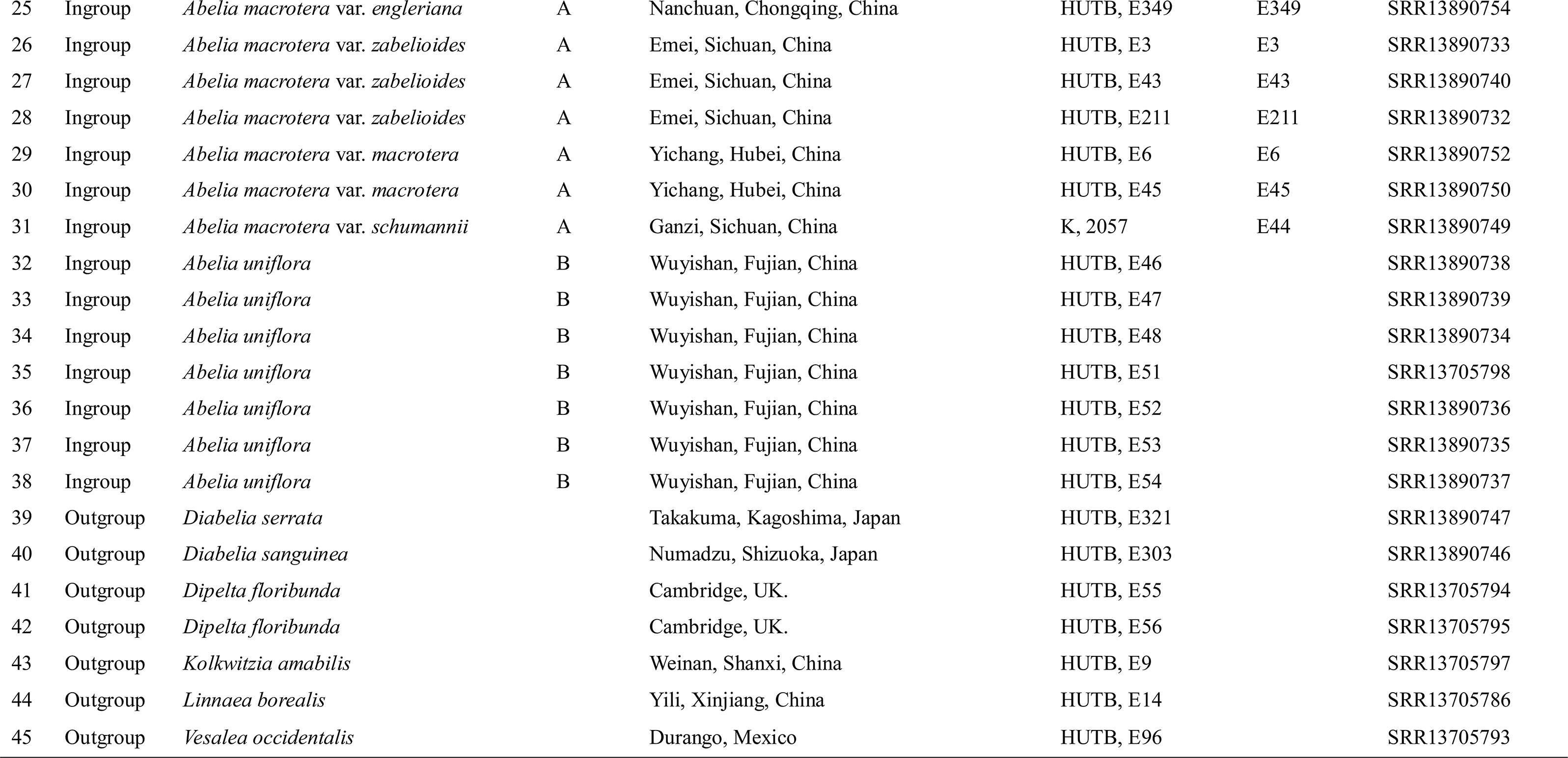
List of species and vouchers used in this study.

### Ecological niche modelling

We performed Ecological Niche Modelling (ENM) on *Abelia* using MAXENT v.3.2.1 (Phillips et al., 2006). For the input we obtained 200 species occurrence data points from the Chinese Virtual Herbarium (CVH, https://www.cvh.ac.cn/), Global Biodiversity Information Facility (http://www.gbif.org/) and field observations. The details of these occurrence data are listed in Table S1. We used 19 bioclimatic variables as environmental data for ENM representing five different time periods. Past and current data for these variables were available from the WorldClim database (http://worldclim.org/). The data included baseline climate (BioClim layers for the period 1950-2000 at a spatial resolution of 30 s arc), data for the last glacial maximum (LGM; ∼22 000 years BP) with spatial resolution at 2.5 min arc resolution simulated by Community Climate System Model (CCSM), Max Planck Institute Model (MPI) and Model for Interdisciplinary Research on Climate (MIROC), and the last interglacial period. Replicate runs (25) of MAXENT were performed to ensure reliable results. Model performance was assessed using the area under the curve (AUC) of the receiver operating characteristic. AUC scores may range between 0.5 (randomness) and 1 (exact match), with those above 0.9 indicating good performance of the model (Swets, 1988). The MAXENT jackknife analysis was used to evaluate the relative importance of the 19 bioclimatic variables employed. All ENM predictions were visualized in ArcGIS v.10.2 (ESRI, Redlands, CA, USA).

### Data accessibility

Raw Illumina data from target enrichment is available at the Sequence Read Archive (SRA) under accession PRJNA706913 (see Table 1 for individual sample SRA accession numbers). DNA alignments, phylogenetic trees and results from all analyses and datasets can be found in the Dryad data repository.

## RESULTS

### Exon assembly

The number of assembled exons per sample ranged from 253 (*A. macrotera* var. *macrotera* E45) to 992 (*A. uniflora* E46) (with > 75% of the target length), with an average of 851 exons per sample (Table S2; Fig. S1). The number of exons with paralog warnings ranged from six in *A. macrotera* var. *macrotera* E45 to 698 in *A. uniflora* E46 (Table S2). The concatenated alignment had a length of 365,330 bp, including 22,045 parsimony-informative sites, and a matrix occupancy of 83% (Table 2). The smallest locus had a length of 150 bp, while the largest locus had a length of 3,998 bp. The plastid alignment resulted in a matrix of 163,130 bp with 3,850 parsimony-informative sites and a matrix occupancy of 96% (Table 2).

**Table 2.**
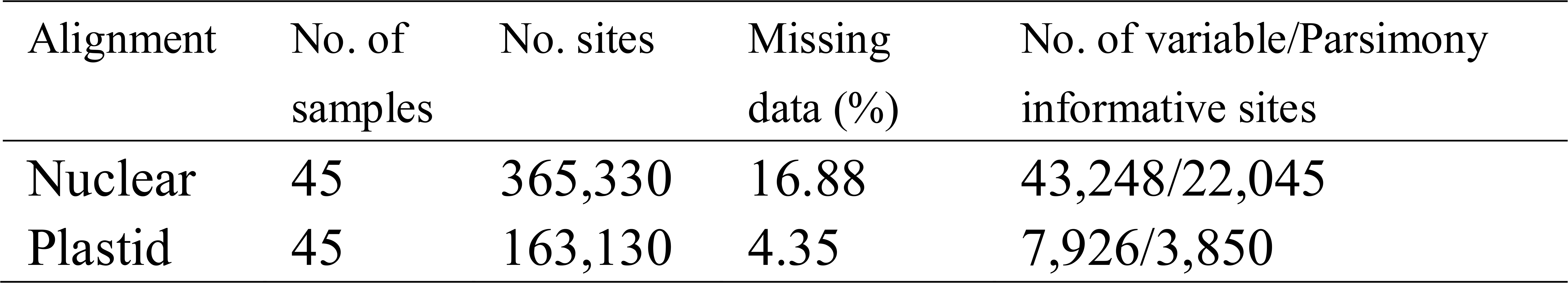
Dataset statistics, including the number of samples, number of characters, number of PI characters, missing data.

### Phylogenetic reconstruction

We inferred the phylogeny of *Abelia* based on nuclear and plastid alignments (Figs. 2-3 and S2-S3). Despite the generally well-supported topology, the positions of clades representing *A. × grandiflora*, *A. uniflora* and *A. macrotera* are unstable across the different phylogenetic analyses. Phylogenetic reconstruction based on the concatenated nuclear data showed all of the major clades are strongly supported (BS ≥75%; Fig. 2, Left). *Abelia chinensis* forms a clade with *A.* × *grandiflora*; and *A. macrotera* var*. zabelioides* is sister to *A. uniflora* with the clade having strong QS support but also a signal of a possible alternative topology (0.97/0/1, Fig. S2, Left). The *A. chinensis + A.* × *grandiflora* clade is then sister to the *A. macrotera* var*. zabelioides + A. uniflora* clade (Fig. 2, Left).

**Figure 2.**
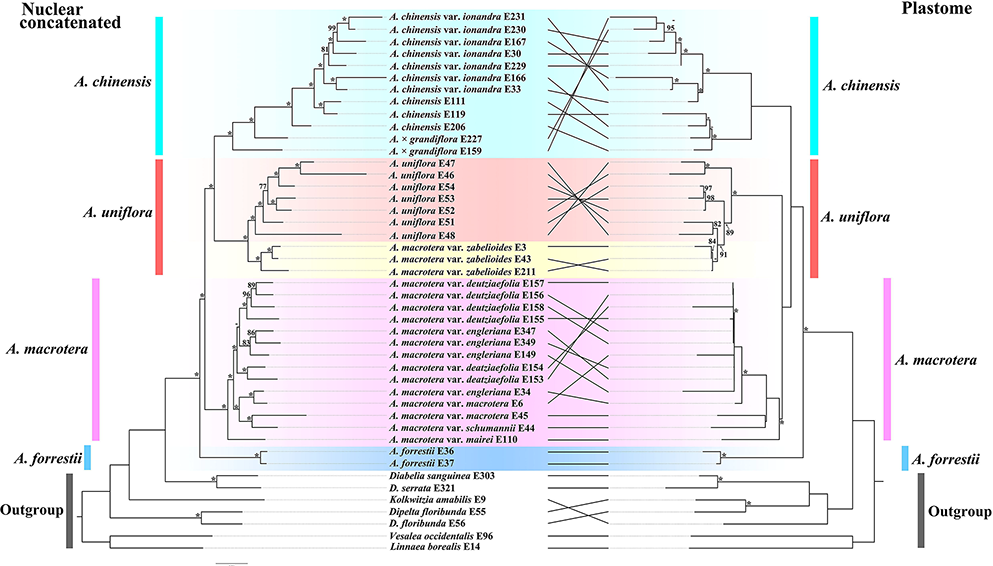
Tanglegram of the nuclear concatenated (Left) and plastid (Right) phylogenies. Dotted lines connect taxa between the two phylogenies. The support values are shown above branches. The asterisks indicate maximum likelihood bootstrap support of 100%. Major taxonomic groups or main clades in the family as currently recognized are indicated by branch colors as a visual reference to relationships.

**Figure 3.**
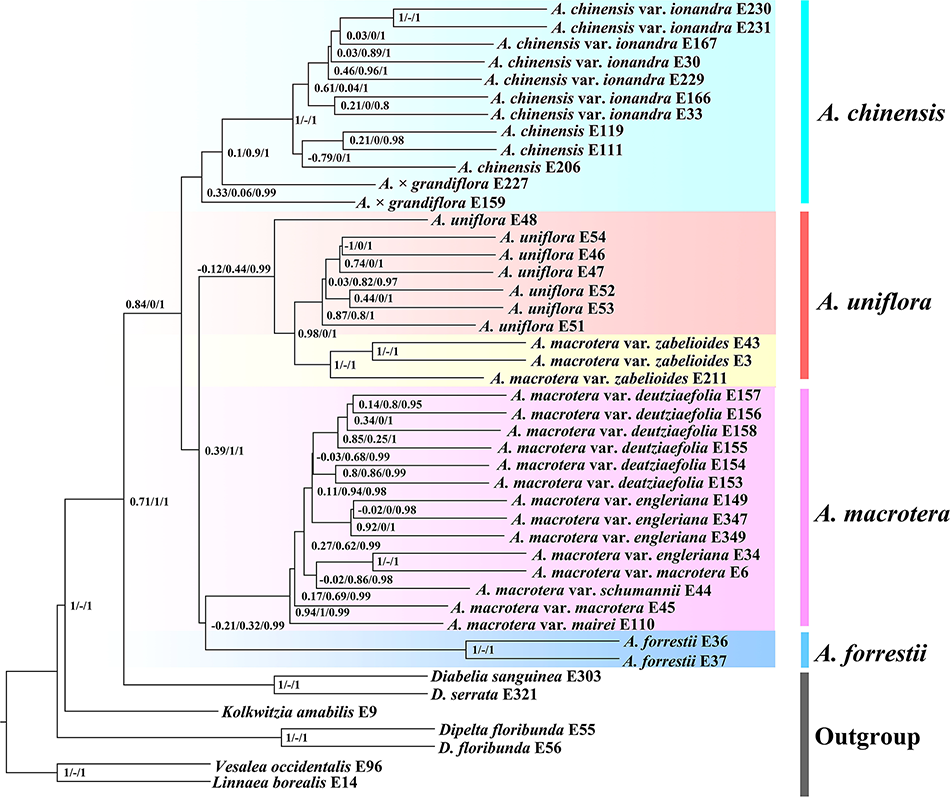
ASTRAL-III species tree; Clade support is depicted as: Quartet Concordance (QC)/Quartet Differential (QD)/Quarte Informativeness (QI)/Local posterior probabilities (LPP).

The ASTRAL analyses (Fig. 3) recovered all major clades with strong support (local posterior probabilities, LPP≥0.95). *Abelia forrestii* is sister to the *A. macrotera* clade, which is then sister to the *A. macrotera* var*. zabelioides + A. uniflora* clade. The *A. chinensis + A.* × *grandiflora* clade is sister to the clade of all remaining species. The clade composed of *A. chinensis + A.* × *grandiflora* has moderate QS support for a signal of a possible alternative topology (0.33/0.06/0.99). The clade composed of *A. macrotera* var*. zabelioides + A. uniflora* has strong QS support but a signal of a possible alternative topology (0.98/0/1). The sister relationship of the *A. macrotera* var*. zabelioides + A. uniflora* has QS counter support as a clear signal for alternative topology (-0.21/0.32/0.99).

For the plastid tree (Fig. 2, Right and S3), *A. macrotera* var*. zabelioides* is nested within *A. uniflora*, with the clade having strong QS support signaling an alternative topology (0.51/0.35/0.98). The *A. chinensis + A.* × *grandiflora* clade shows a sister relationship with the *A. macrotera* var*. zabelioides + A. uniflora* + *A. macrotera* clade, with strong QS support (0.75/0.51/0.98). The results strongly support *A. forrestii* from Yunnan, China as sister to the remainder of the genus with the QS analysis providing full support (1/–/1, Fig. S3).

In our species network analyses (including both 15 and 12 samples), all networks with one to five reticulation events were shown to be a better model than a strictly bifurcating tree (Table 3; Figs. 4 and S4-S5). Four reticulation events were inferred for the best network in the 15 samples tree, two reticulation-events involving *A. × grandiflora*, the third reticulation event occurred to *A. chinensis* and the fourth revealed ancestral gene flow to the *A. × grandiflora* clade (Fig. 4A). For *A. × grandiflora* E159, the inheritance probability analysis showed that the major genetic contributor (57.2%) came from the ancestor of *A. × grandiflora* and a minor contribution was from the *A. macrotera* clade (42.8%). For *A. × grandiflora* E227, the major genetic contributor came from *A. uniflora* (53.6%). For *A. chinensis* E206, the major genetic contributor (53.0%) came from the ancestor of *A. × grandiflora*. For the ancestor of the *A. × grandiflora* clade, the major genetic contributor was a putative ancestor of *A. chinensis* (74.1%) with a minor contribution from *A. uniflora* (25.9%). In the 12 samples tree (Fig. 4B), we removed the known hybrids from the previous analyses by excluding *A.* × *grandiflora* E159, *A.*× *grandiflora* E227 and *A. chinensis* E206, and then discovered that the best network had three reticulation events. There appears to be abundant gene flow in the backbone of *Abelia* and the 12 samples analyses showed that *A. macrotera*, *A. uniflora* and *A. chinensis* were each derived from complex hybridization events.

**Figure 4.**
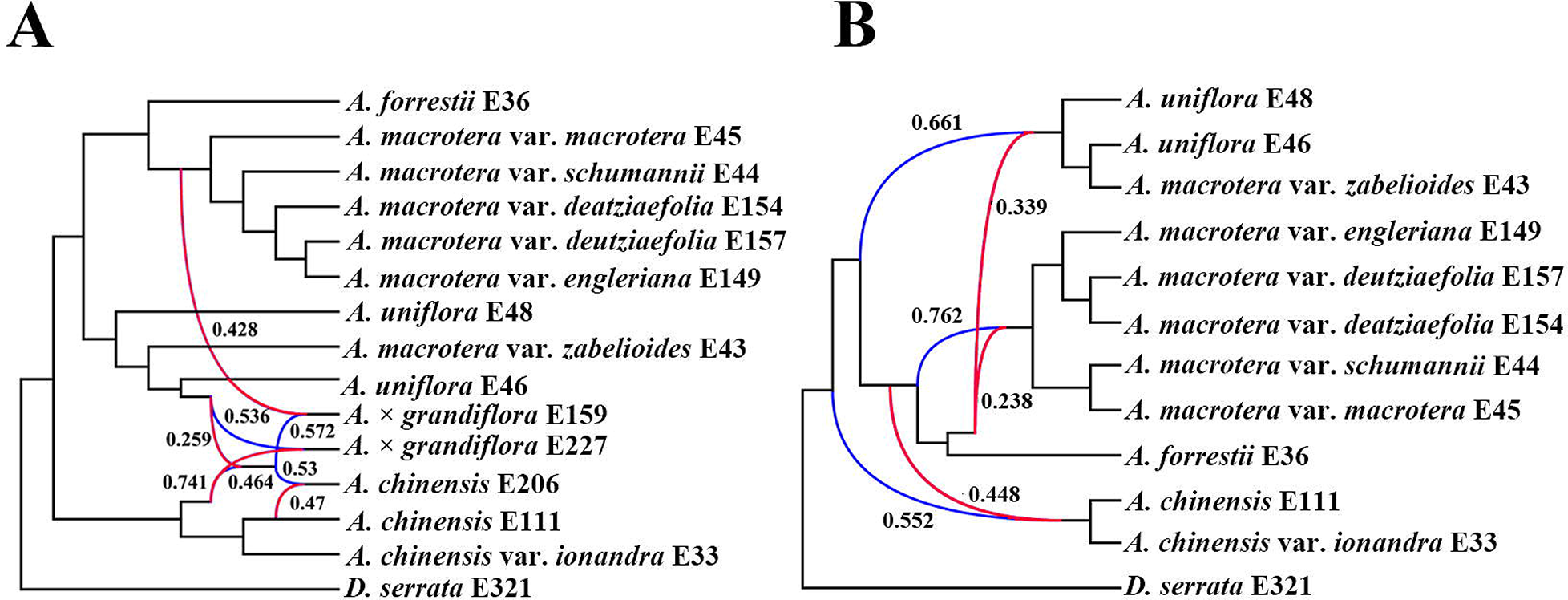
Best supported species networks inferred with PhyloNet for the (A) 15 samples, (B) 12 samples. Numbers next to the inferred hybrid branches indicate inheritance probabilities. Red lines represent minor hybrid edges (edges with an inheritance contribution < 0.50).

**Table 3.**
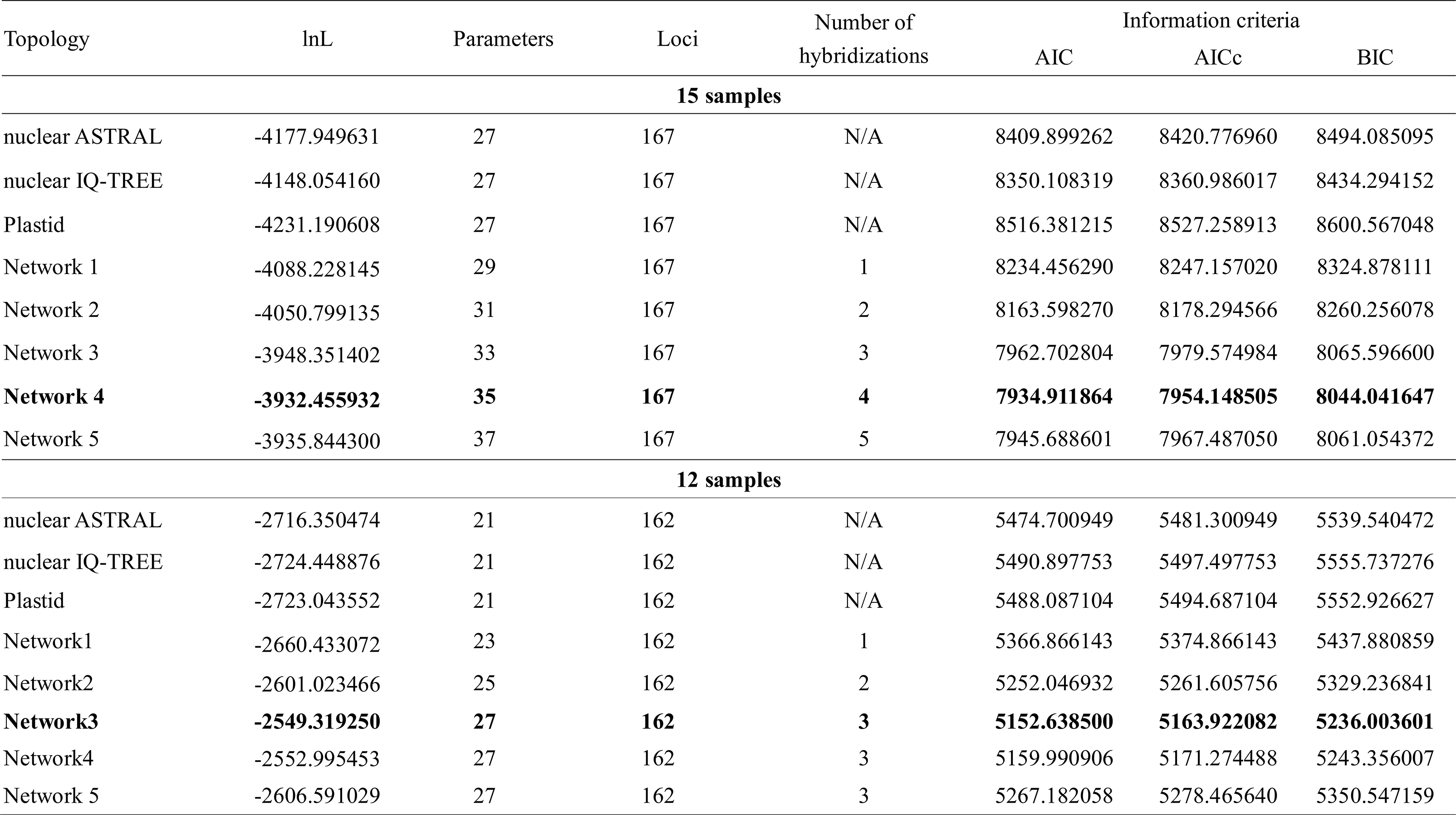
Model selection of the different species networks and bifurcating trees.

### Divergence time estimation

Divergence times were estimated using BEAST v.1.8 which takes into account phylogenetic and calibration uncertainties. Given that *A. × grandiflora* is an artificial hybrid, the taxon should not be included in the divergence time estimation and ancestral area reconstruction analysis of the genus.

Divergence time estimates based on the nuclear tree (Fig. 5) suggested that the ancestor of *Abelia* originated 33.47 Ma (95% HPD: 23.53-38.21 Ma, node 1). The divergence of *A. chinensis* clade from the remaining *Abelia* species could be dated to 29.56 Ma (95% HPD: 21.64-33.76 Ma, node 2). *Abelia forrestii* diverged from its close relatives 24.93 Ma (95% HPD: 18.62-29.16; node 3). The split between *A. uniflora* and *A. macrotera* var*. zabelioides* was 14.93 Ma (95% HPD: 12.39-20.21 Ma, node 6). *Abelia macrotera* diverged from its close relatives 23.51 Ma (95% HPD: 16.91-27.09; node 4). *Abelia chinensis* var*. ionandra* diverged from its close relatives 12.44 Ma (95% HPD: 8.61-17.03 Ma, node 7). A comparison of the divergence time estimates using the chloroplast tree is shown in Fig. S6. For instance, the age of the stem group of *Abelia* was dated to 45.10 Ma (95% HPD: 34.91-50.09 Ma, node 1). *Abelia forrestii* diverged from its close relatives 29.75 Ma (95% HPD: 21.64-39.04; node 2). The split between *A. uniflora* and *A. macrotera* var*. zabelioides* was 14.31 Ma (95% HPD: 8.16-23.08 Ma, node 6).

**Figure 5.**
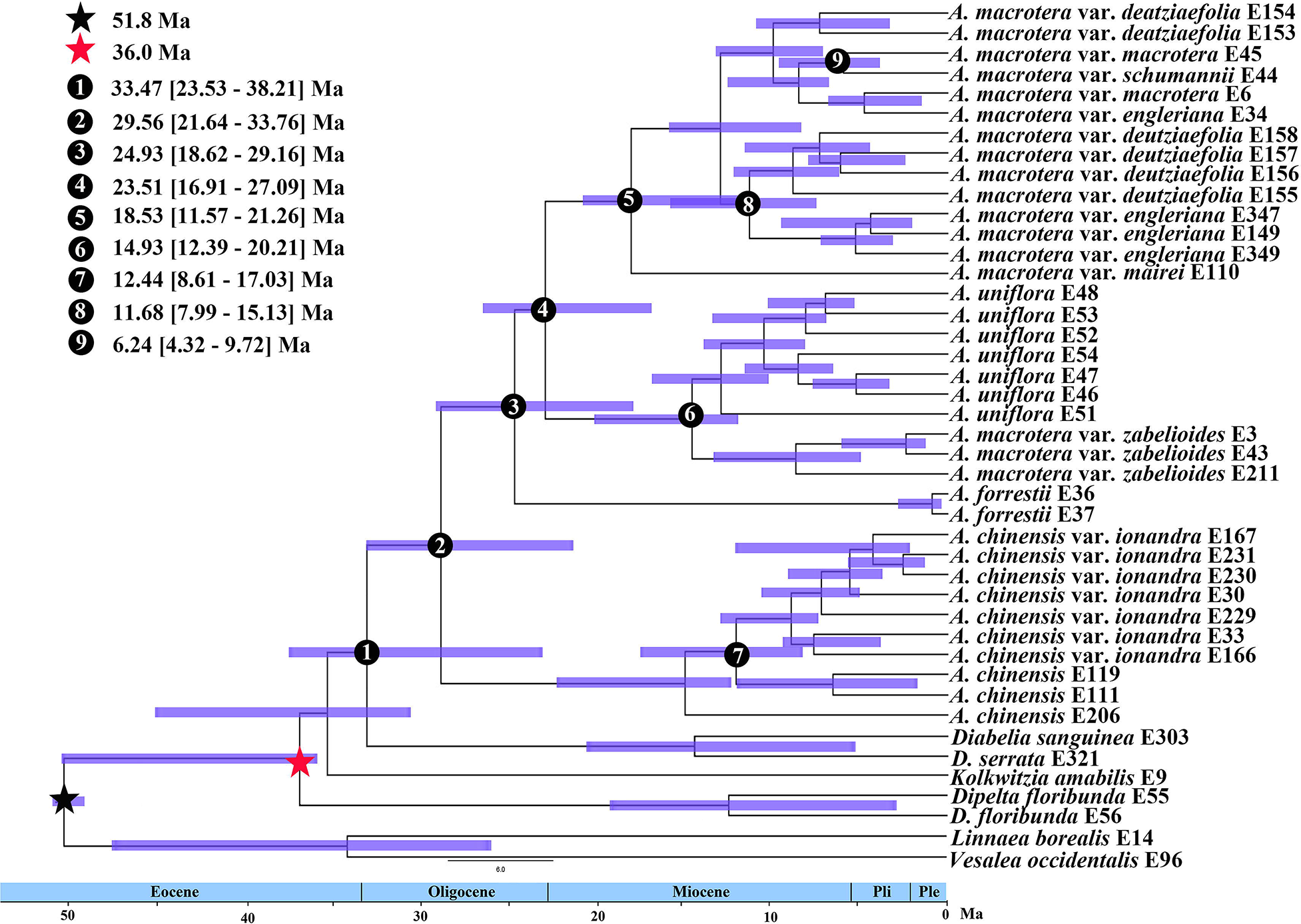
BEAST analysis of divergence times based on nuclear data. Calibration points are indicated by stars. Numbers 1-9 represent major divergence events in *Abelia*; Mean divergence times and 95% highest posterior densities are provided for each.

### Ancestral area reconstruction

According to our ancestral area reconstructions paired with divergence time estimation, *Abelia* originated around the early Eocene (Fig. 5) in mainland China (Fig. 6). Our analyses revealed two dispersal events and three vicariance events among the three defined biogeographic areas (Fig. 6). The ancestor of *Abelia* was inferred to have been present throughout region A (Yunnan, Chongqing, Sichuan, Guizhou and Hubei) and B (Jiangxi and Fujian), with subsequent dispersal or vicariance across southeastern China. The results indicate a southwestern and central-eastern China origin for the genus *Abelia*.

**Figure 6.**
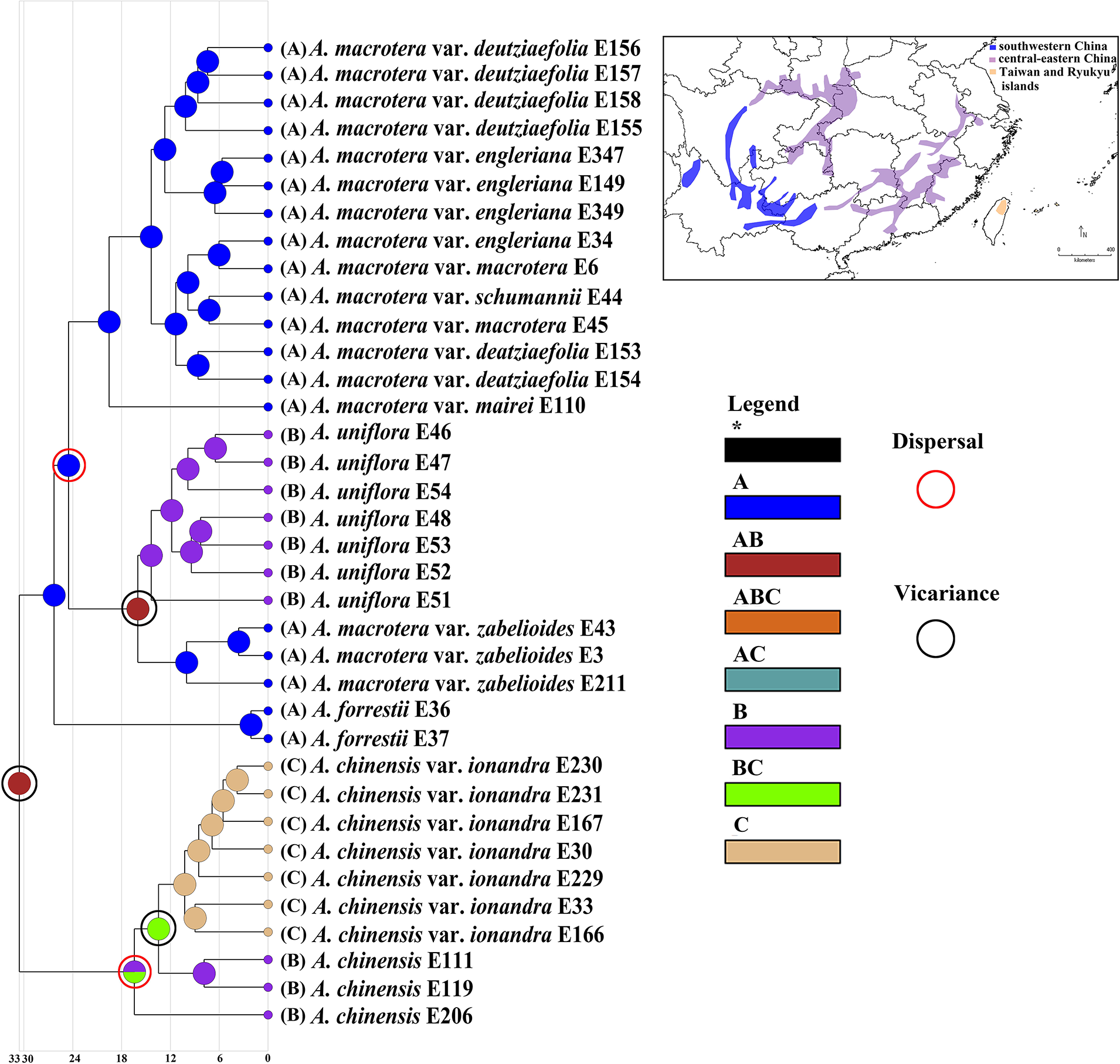
Ancestral area reconstruction for *Abelia*. Areas of endemism are as follows: (A) southwestern China, (B) central-eastern China, (C) Taiwan and Ryukyu islands. The numbered nodes represent crown nodes of important colonization events.

### Ecological niche modelling and niche identity tests

The AUC test result (mean ± SD) for the ENM, averaged across all 25 runs, was significant (0.996 ±0.000 for *A. chinensis* and 0.993± 0.001 for *A. uniflora*) (Figs. 7 and S7), which supports the predictive power of the model. The MaxEnt analyses show that the current distribution prediction is largely identical to the recorded distribution of *Abelia*, and expansion beyond the existing range is also reasonable, including *A. uniflora* populations along the narrow coastline of eastern China and Japan (Fig. S7). In our analysis, the LGM climate and the present time climate species distribution areas show different distribution patterns. We used CCSM, MIROC and MPI models to simulate the LGM climate for the five species of *Abelia*, and the inferred distribution from the two LGM models (CCSM and MPI models) were highly similar. For the CCSM and MIROC models (the MIROC model area simulation during the LGM period is warmer and drier than the CCSM model), the distribution density and range of *Abelia* under the MPI model was slightly lower than the those generated from the CCSM model, but the differences were not significant (Fig. S7). Compared with the LGM climate, the potential distribution range of *Abelia* species in mainland China will be much smaller, while the habitat remains stable in most of the present time climate distributions.

**Figure 7.**
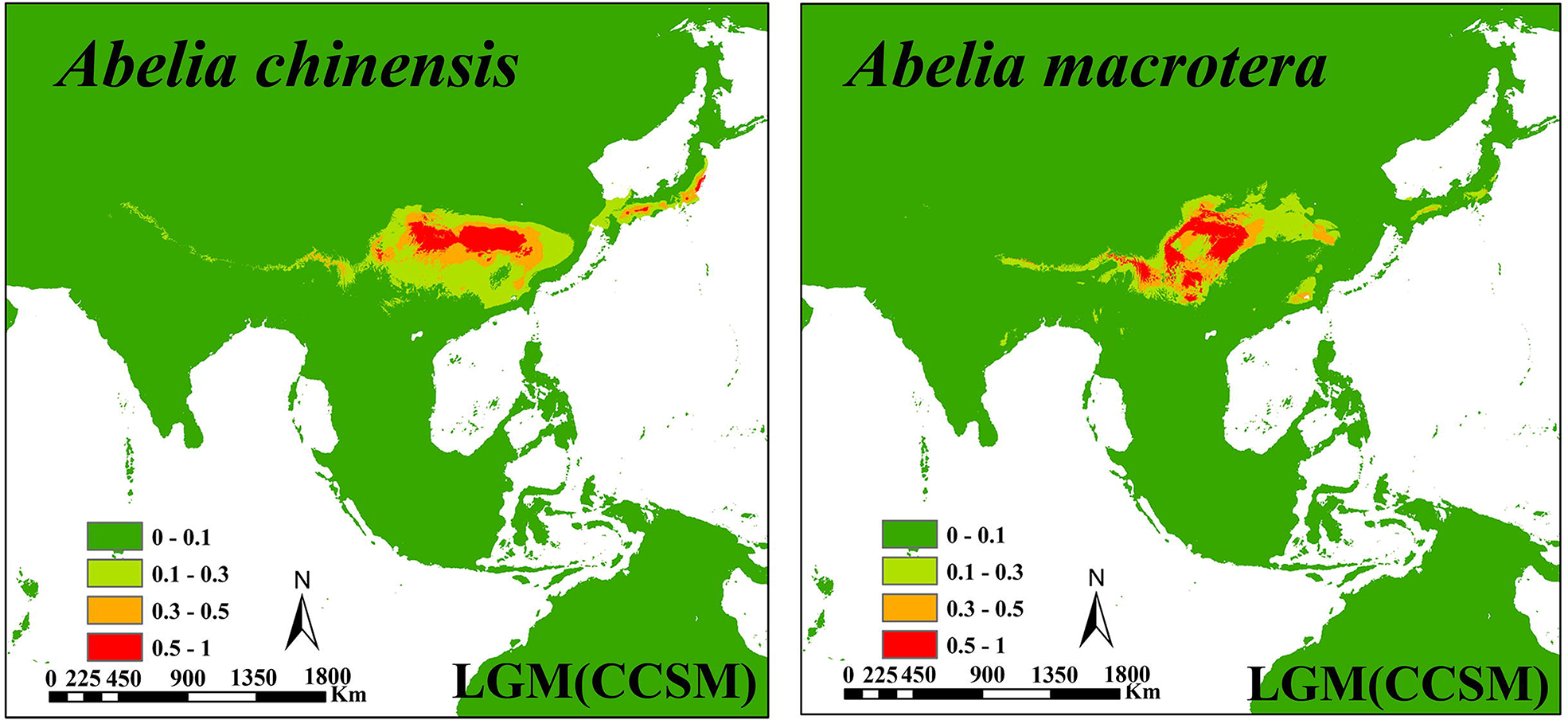
Comparison of potential distributions as probability of occurrence for *A. chinensis*, and *A. uniflora* at the CCSM climatic scenarios of the Last Glacial Maximum (LGM, ca. 21,000 years BP). The maximum training sensitivity plus specificity logistic threshold has been used to discriminate between suitable (cutline 0.1-1 areas) and unsuitable habitat. The darker color indicates a higher probability of occurrence.

## DISCUSSION

### Phylogenetic relationships within Abelia

*Abelia* is a genus in the Sino-Japanese Floristic Region with a disjunct distribution between the Himalaya-Hengduan Mountains and Taiwan (Kim et al., 1998; Wang et al., 2015). The genus was established by R. Brown in 1818 with *A. chinensis* R. Brown as the type species. Based on morphological features, *Abelia* was considered to be closely related to *Linnaea* (Rehder, 1911). Hu (1988) divided *Abelia* into two genera: *Abelia* s.s. and *Zabelia* (Rehd.) Makino. Later, Kim (1998) found that *Abelia* was sister to *Diabelia* based on a number of traditionally used morphological characters (mostly inflorescence traits) and molecular data. Previous studies did not resolve relationships within *Abelia*, likely because of the limited informative sites in the loci used (Kim, 1998; Jacobs et al., 2010; Wang et al., 2015; Landrein, 2017).

This study resolved well-supported phylogenetic relationships within *Abelia* using both nuclear and plastid data (Figs. 2-3 and S2-S3). The inferred topologies of the four recognized species in the nuclear and plastid trees are not entirely congruent (Figs. 2-3 and S2-S3). For instance, in the nuclear concatented tree (Fig. 2, Left), *Abelia macrotera* var. *zabelioides* is sister toand the *A. uniflora* form a clade, with the nodes having strong QS support values (Fig. 3). In the plastid tree (Figs. 2, Right) and the ASTRAL tree (Fig. 3), *Abelia macrotera* var*. zabelioides* is nested within *A. uniflora.* Cytonuclear discordance has been widely detected in plants (Lee et al., 2019; García et al., 2017; Hodel et al., 2021; Wang et al., 2021). The three main reasons for cytonuclear discordance are gene duplication/extinction, horizontal gene transfer/hybridization, and incomplete lineage sorting (ILS), but sources of such discordance remain hard to disentangle, especially when multiple processes co-occur (Degnan and Rosenberg, 2009).

*Abelia macrotera* var. *zabelioides* and *A. uniflora* share a recent common ancestor (Landrein et al., 2017), which is consistent with the results of our phylogenetic analysis (Figs. 2-3 and S2-S3). The species network analysis (Fig. 4) showed the two species are formed by different hybridization events.

### Hybridization and introgression

There are controversial taxa or species complexes in *Abelia.* The *A. uniflora* complex is the most enigmatic in the genus, though it was only briefly described by Wallich (1829). Members of the *A. uniflora* complex are often difficult to identify with morphological characters alone. The morphological differences among entities within the *A. uniflora* complex can be explained by one or several historical or ongoing evolutionary processes. Specifically, the null taxonomic hypothesis is that the *A. uniflora* complex is a single hyperdiverse species (Link-Pérez, 2010), while the alternative is that speciation has occurred or is underway. We expect to detect clear genetic differentiation, which may be stronger if speciation is complete, or weaker when speciation is ongoing or confounded by introgression (Nosil, 2008; Liu et al., 2018). Based on morphological evidence and AFLP data, Landrein et al. (2017) indicated that *A. uniflora* shows evidence of formation due to hybridization/ introgression between *A. macrotera* and *A. chinensis*. Additionally, Landrein et al. (2017) showed that *A. forrestii* occurs in the contact zone between *A. macrotera* and *A. chinensis*, leading to a potential hypothesis of hybrid/introgression origin for *A. forrestii* from *A. macrotera* and *A. chinensis*.

In our analysis, it was found that there may be widespread cytonuclear discordance exists across *Abelia* (Figs. 2-3 and S2-S3). The discordance of the phylogeny may be caused by ILS or hybridization. Hybridization, introgression, and horizontal gene transfer are all examples of gene flow across populations that may be modeled by phylogenetic networks. Sols-Lemus et al. (2017) proposed that a minor contribution (∼ 0.10) from a parental population to a reticulate node may indicate an introgression event based on inheritance probabilities. Inheritance probabilities near 0.50 may indicate that the reticulate node is the result of hybrid speciation between the parental populations. Our results show that the parental contributions to the reticulation events detected in *Abelia* (Fig. 4A) are essentially equal. For *A. × grandiflora* E159, the inheritance contributions (0.428 and 0.572) support a hybridization event between the ancestral lineage of the *A. macrotera* clade and *A. chinensis* E206. For *A. × grandiflora* E227, the inheritance contributions (0.536 and 0.464) support a hybridization event between *A. uniflora* and the ancestral lineage of the *A. chinensis* clade. For *A. chinensis* E206, the inheritance contributions (0.572 and 0.470) support a hybridization event between *A. × grandiflora* E227 and *A. chinensis* E111. For the ancestor of the *A. × grandiflora* clade, the reticulation event reveals that there has been extensive gene flow between the *A. chinensis* clade and *A. uniflora*. However, *A. uniflora* has less genetic contribution (0.259), suggesting that ancestral gene flow from *A. uniflora* to the ancestor lineage of the *A. × grandiflora* clade could also produce the observed results (Fig. 4A). Because inheritance probability can be influenced by many biological factors, additional biological information is needed to interpret these values reliably (Sols-Lemus et al., 2017). In the 12 samples tree (Fig. 4B), we discovered that the ancestral lineage of *A. macrotera*, *A. uniflora* and *A. chinensis* were each derived from complex hybrid origins, which can explain the complicated taxonomy of the group.

### Diversification history of Abelia

In this study, the dating results are incongruent between the nuclear and plastid analyses, with the nodes in the plastid tree generally showing older dates than those in the nuclear tree. Generally, chloroplast genes have shortcomings such as low selection pressure, single-parent inheritance, and slow inheritance rate (Lanier and Knowles, 2012; Wang and Liu, 2016), therefore we chose nuclear data for our dating analysis. Our analyses indicate that the stem of *Abelia* dates to the early Eocene 33.47 Ma (95% HPD: 23.53-38.21 Ma) (Fig. 5). Most of the diversification events within *Abelia* were dated to the early to middle Miocene (Fig. 5).

Our phylogenetic analyses indicate that *A. chinensis* var. *ionadra* from the island of Taiwan is sister to *A. chinensis* from Jiangxi province, China (Figs. 2-3 and S2-S3). In addition, previous phylogenetic and phylogeographic studies have generally shown that existing populations of many species on the island of Taiwan are descendants of mainland China (e.g. in *Taiwania*, Chou et al., 2011; *Sassafras*, Nie et al., 2007; *Dichocarpum arisanense*, Xiang et al., 2017; *Triplostegia*, Niu et al., 2018), with few reports supporting a reverse scenario. Based on our divergence time estimates, the stem age of *A. chinensis* var. *ionadra* was dated to the middle Miocene, ca. 12.44 Ma (Fig. 5, node 7), which greatly predates the formation of the island of Taiwan, having originated in the late Miocene (Sibuet and Hsu, 2004). A similar pattern was shown in *Tinospora dentata* which split from its Japanese relatives in the early Miocene around 38.97 Ma (95% HPD: 26.45-52.52) (Wang et al., 2017).

The island of Taiwan is located at the eastern edge of the Eurasian plate and the Taiwan Strait could have first occurred in the late Mesozoic (Ye, 1982; Suo et al., 2015). Owing to the intense tectonic movements, contacts between the island of Taiwan and mainland China occurred during the late Cretaceous-Early Paleocene and late Eocene-early Oligocene (Teng, 1992). One hypothesis that has been put forward is that tropical rainforests in the Indo-Malayan region may have appeared near the Cretaceous-Paleogene (K-Pg) boundary (Wang et al., 2012). Plant macrofossils that are closely related to extant subtropical species have been recorded from Taiwan during the early Miocene, while pollen fossils date to the Oligocene (Li, 2000). Due to the landmass being subjected to stretching and rifting (Ye, 1982; Teng, 1992; Suo et al., 2015), part of the mountain ranges on the island of Taiwan represents a relict area of the ‘Taiwan-Sinzi Folded Zone’ (Ye, 1982; Teng, 1992; Li and Wen, 2016), which facilitated a cooler climate in Asia during the late Eocene (Zachos et al., 2001). Around this time, a vicariance event between Taiwan and mainland China may have occurred. Based on the results of our *Abelia* sequencing, the Taiwan (Yilan) flora may contain relatively ancient components (Li and Wen, 2016). The flora of mainland China has been acknowledged to have contributed to Taiwan’s floristic assembly (Li and Keng, 1950). The Luzon arc did not begin to collide with the Eurasian plate until the late Miocene (9-6 Ma) (Sibuet et al., 2002). Thus, we hypothesize that the relatively young gene flow in the flora of Taiwan may be derived from plants in mainland China. This hypothesis needs further validation by combining phylogenetic and molecular dating methods, as well as by studying other plant groups, particularly more ancient lineages.

The stem group of *Abelia* is dated to have formed during the early Eocene (33.47 Ma, 23.53-38.21 Ma; node 1), later than the early uplift of the Qinghai-Tibet Plateau (QTP). However, ancestral area reconstructions revealed that the *A. uniflora* complex might have colonized southwestern China, central-eastern China, and Taiwan and Ryukyu islands (Fig. 6).

### Potential refugia and demographic history

Many botanists and biogeographers have explained the discontinuous distribution pattern of plants in East Asia and North America from different perspectives (Xiang et al., 1998; Wen, 2001; Nie et al., 2007; Sessa et al., 2012). It is hypothesized that East Asia was less affected by the Quaternary glaciations than North America and Nordic Europe, except in alpine areas (Shi et al., 2006).

Climate shifts in the Quaternary and associated environmental changes may have driven the diversification of *Abelia*. Diversification would have been sustained by Pleistocene glaciations because species habitats underwent expansion and contraction, leading to populations flourishing, migrating or extincting (Comes et al., 2008; Qiu et al., 2011).

Our ENM analysis provides a detailed hypothesis during the last glacial cycle (Figs. 7 and S7). The results suggest that extant populations likely differentiated well before the LGM and also provide strong evidence for survival of *Abelia* in southwest China during the LGM (Fig. S8). Therefore, the climatic shock during the LGM may have had a significant impact on the recent history of *Abelia* at the population level compared with an earlier period of species differentiation (Qiu et al., 2011). Potential habitat for *A. forrestii* spanned most of southwest China during the LGM period. However, its current distribution is only in Yunnan and Sichuan of southwestern China from 1900 to 3300 m in altitude. *Abelia chinensis* and *A. uniflora* were distributed in mainland China and Southern Japan during the LGM period (Fig. 7). However, the current distribution area has shrunk, which highlights the importance of topographic complexity in promoting species differentiation both by increasing habitat diversity and limiting gene flow between elevation-restricted populations (Ye et al., 2019). Yunnan, Guizhou, Sichuan and Guangxi are placed in the East Asiatic region in terms of phylogenetic regionalization, but are placed in the Paleotropic region for taxonomic regionalization. Because these areas possess more endemism, particularly paleoendemic genera (Ying et al., 1994), they are thought to have been a refuge during the Pleistocene glacial period (López-Pujol and Ren, 2010; Ye et al., 2019). For *A. macrotera*, there is few significant change when comparing its LGM distribution to that in the current period. Thus, we hypothesize that the refugia reflect the evolutionary history of *A. macrotera*. This hypothesis needs to be further tested by studying additional plant groups, especially ancient lineages, through the integration of plastome and adequate nuclear data to balance the preferences towards nuclear/chloroplast genomes. In addition, *Abelia chinensis* var. *ionandra* has expanded to the coastal regions of East China and reached Taiwan via land bridges across the Taiwan Strait during the glacial period. Due to repeated glacial-interglacial cycles, mainland China and Taiwan populations may have experienced frequent gene flow through a continental bridge during Quaternary. After the LGM, *Abelia* populations in southeastern China may have experienced a large-scale extinction event, and Jiangxi may be a glacial refuge for *A. chinensis*. The uplift of the QTP was a consequence of the collision of the Indian subcontinent with the Eurasian plate 40-50 million years ago (Yin and Harrison, 2000; Fu et al., 2020). Because of multiple asynchronous expansions, multiple independent refugia *in situ* and complex topography in this region might explain why accurate analyses of species distribution patterns are difficult (Cun and Wang, 2010; Yan et al., 2012). Because *Abelia* does not have a strong ability of pollen or seed transmission, the present distribution pattern of *A. chinensis* var. *ionandra* may be the result of topographic isolation and postglacial contraction.

## CONCLUSIONS

Natural hybridization/infiltration is extensive in *Abelia* and may require further investigation with extensive populational sampling. Our findings support the use of Hyb-Seq data at the population level to clarify the boundaries among species complexes. For other taxa, our study demonstrates the importance to use both nuclear and plastid genome data, as well as network analysis, to untangle the potential reticulation history. Our results also indicate that several interglacial refuges in *Abelia* have been identified in the middle of the Hengduan Mountains. Besides, our research provide a theoretical foundation for investigating the disjunctively distributed plants between Taiwan island and mainland China, as well as supporting hypotheses of postglacial range contraction together with topographic heterogeneity being responsible for the observed disjunction.

## TAXONOMIC TREATMENT

***Abelia uniflora*** R. Br. in Wall., Pl. As. Rar. 1. 15. 1830. -Type: China, Fujian, [‘Ngan-Ke-Kyen’Black Tea Country], *Reeves s.n.* (holotype: CGE!, isotype: BM!). =*Abelia macrotera* (Graebn. & Buchw.) Rehder var. *zabelioides* Landrein, Kew Bull. 74(4)-70: 93. 2019. Type: China, Sichuan, Leshan Pref., E’mei Shan, 21 June 1938, *H. C. Chow 7573* (holotype: A!). **Synonym nov.**

## Author contributions

H.F.W. and J.W. conceived the study. H.F.W. and D.F.M-B. performed the research and analyzed the data. Q.H.S., D.F.M-B., H.X.W., J.W. J.B.L, and H.F.W. wrote and revised the manuscript.

## ACKNOWLEDGEMENTS

The work was funded by National Science Foundation of China (31660055) and Hainan Provincial Natural Science Foundation of China (421RC486). We appreciate Gabriel Johnson for his help with the target enrichment experiment, and the United States National Herbarium for collection access. We acknowledge the staff in the Laboratories of Analytical Biology at the National Museum of Natural History, the Smithsonian Institution for technical support and assistance.

---

**Fig. S1.**
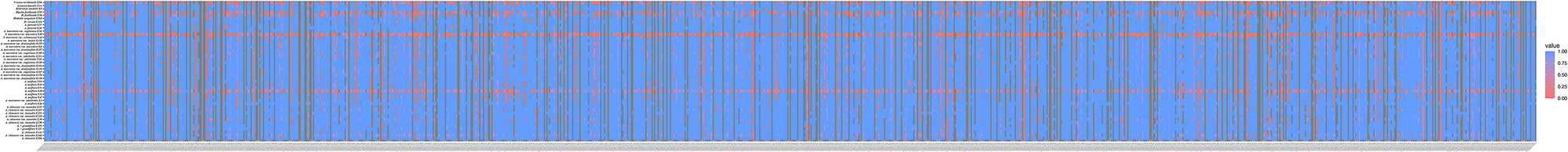
Heatmaps showing gene recovery efficiency for the nuclear gene in 45 species of *Abelia*. Columns represent genes, and each row is one accession. Shading indicates the percentage of the reference locus length coverage.

**Fig. S2.**
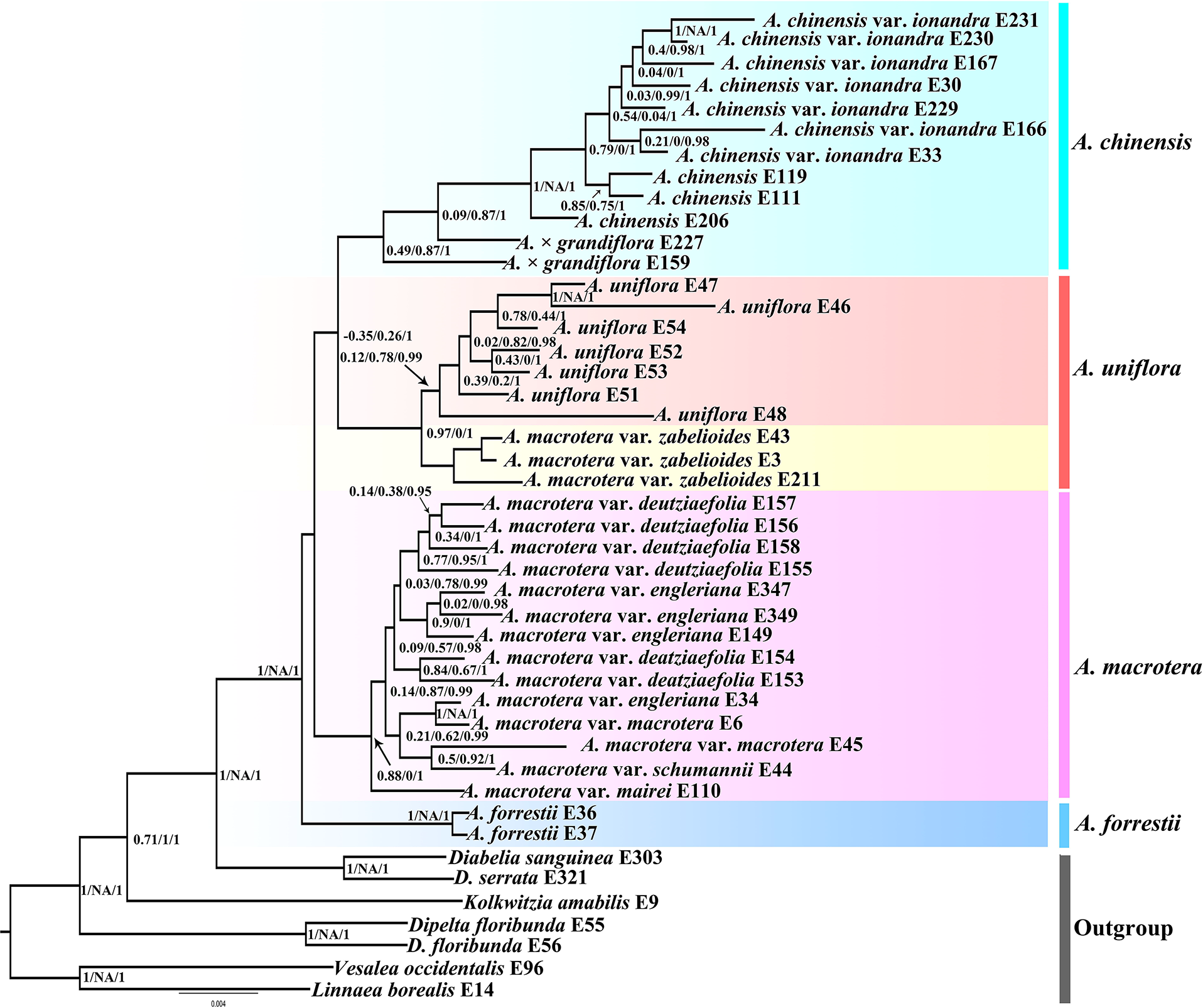
Nuclear concatenated IQ-tree; Clade support is depicted as: Quartet Concordance (QC)/Quartet Differential (QD)/Quarte Informativeness (QI).

**Fig. S3.**
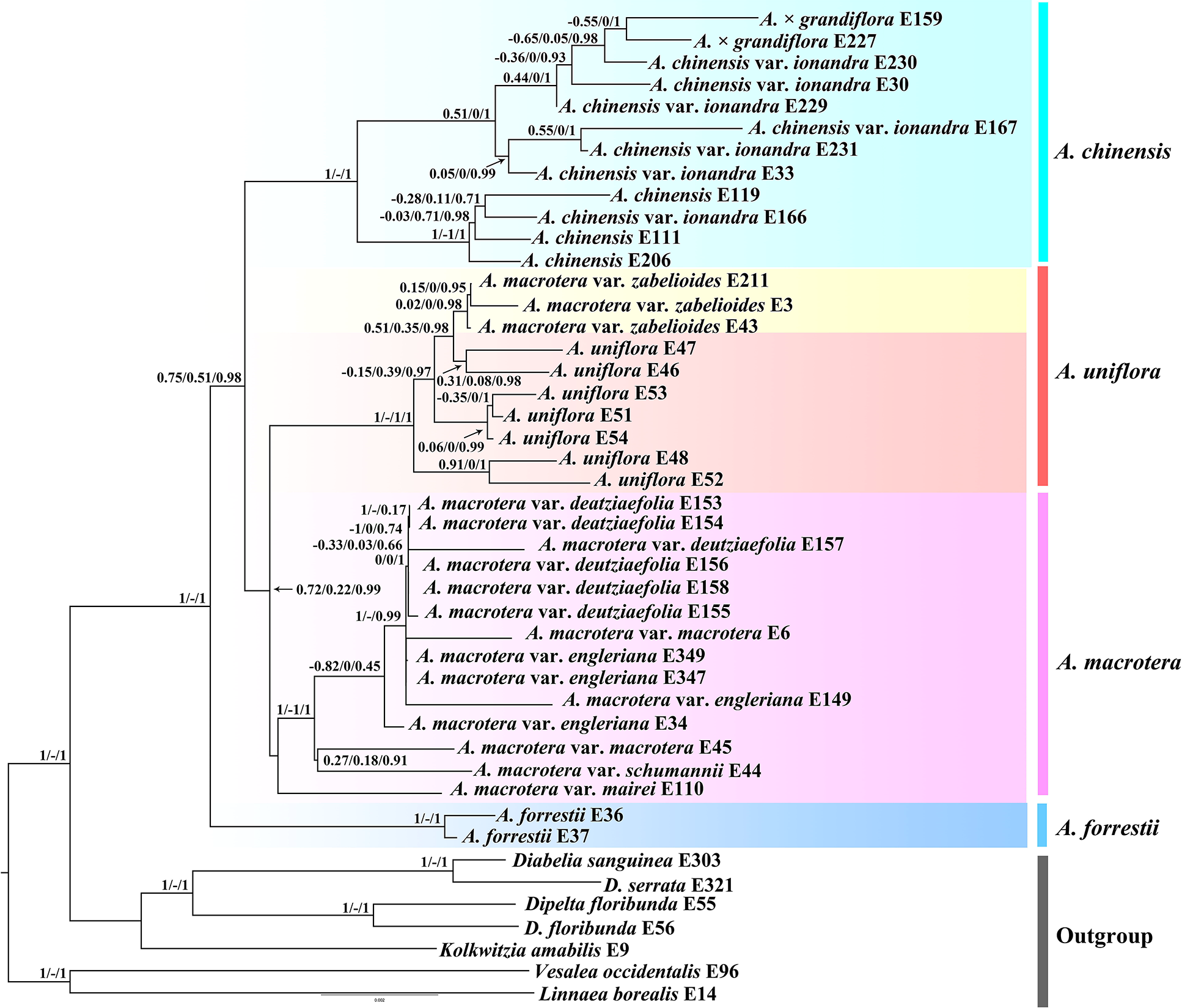
Plastid IQ-tree; Clade support is depicted as: Quartet Concordance (QC)/Quartet Differential (QD)/Quarte Informativeness (QI)/Local posterior probabilities (LPP).

**Fig. S4.**
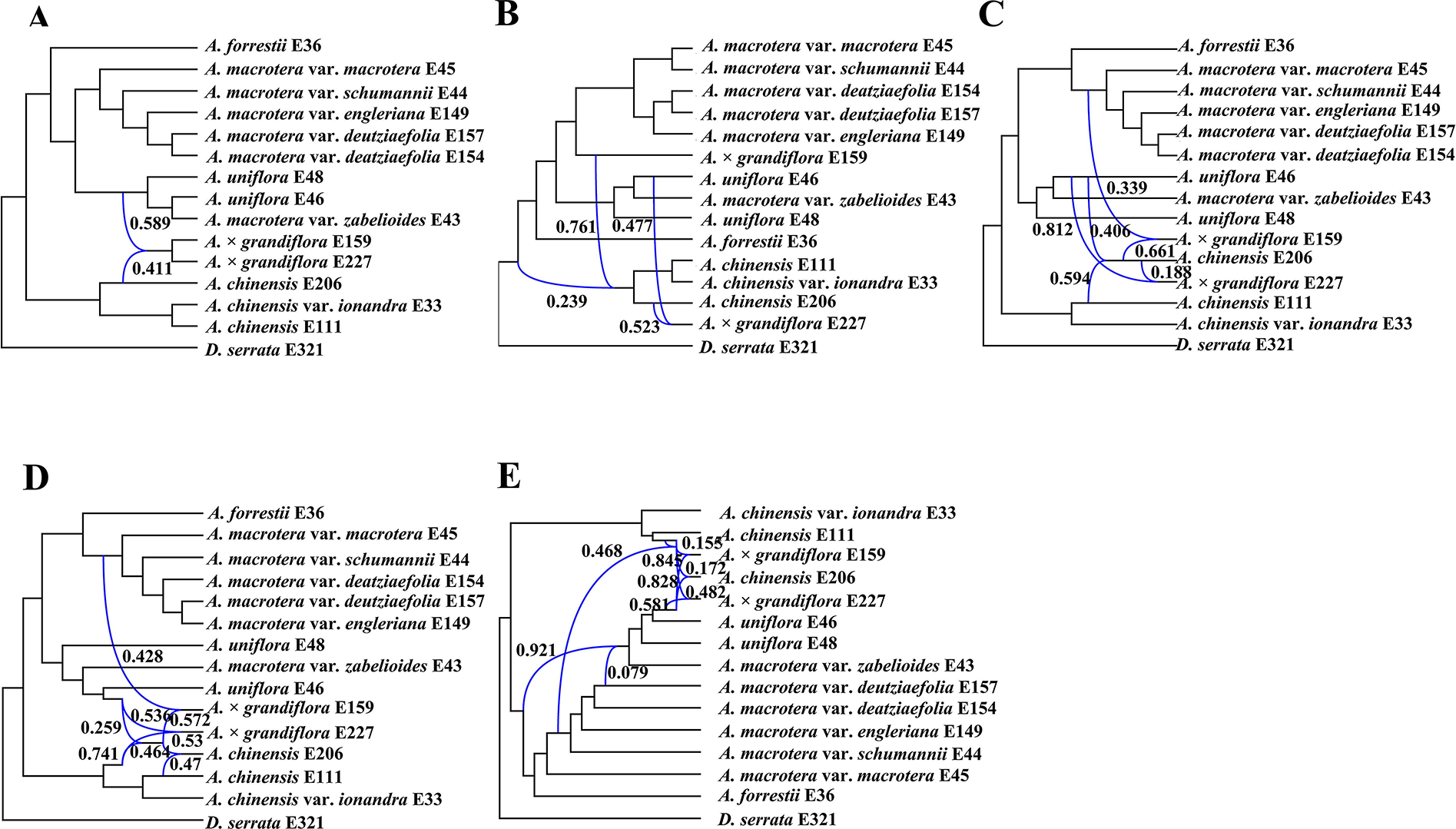
Best species networks of the selective nuclear data estimated with PhyloNet for the 15 samples data set. A: One hybridization event; B: Two hybridization events; C: Three hybridization events; D: Four hybridization events; E: Five hybridization events. Blue branches connect the hybrid nodes. Numbers next to blue branches indicate inheritance probabilities.

**Fig. S5.**
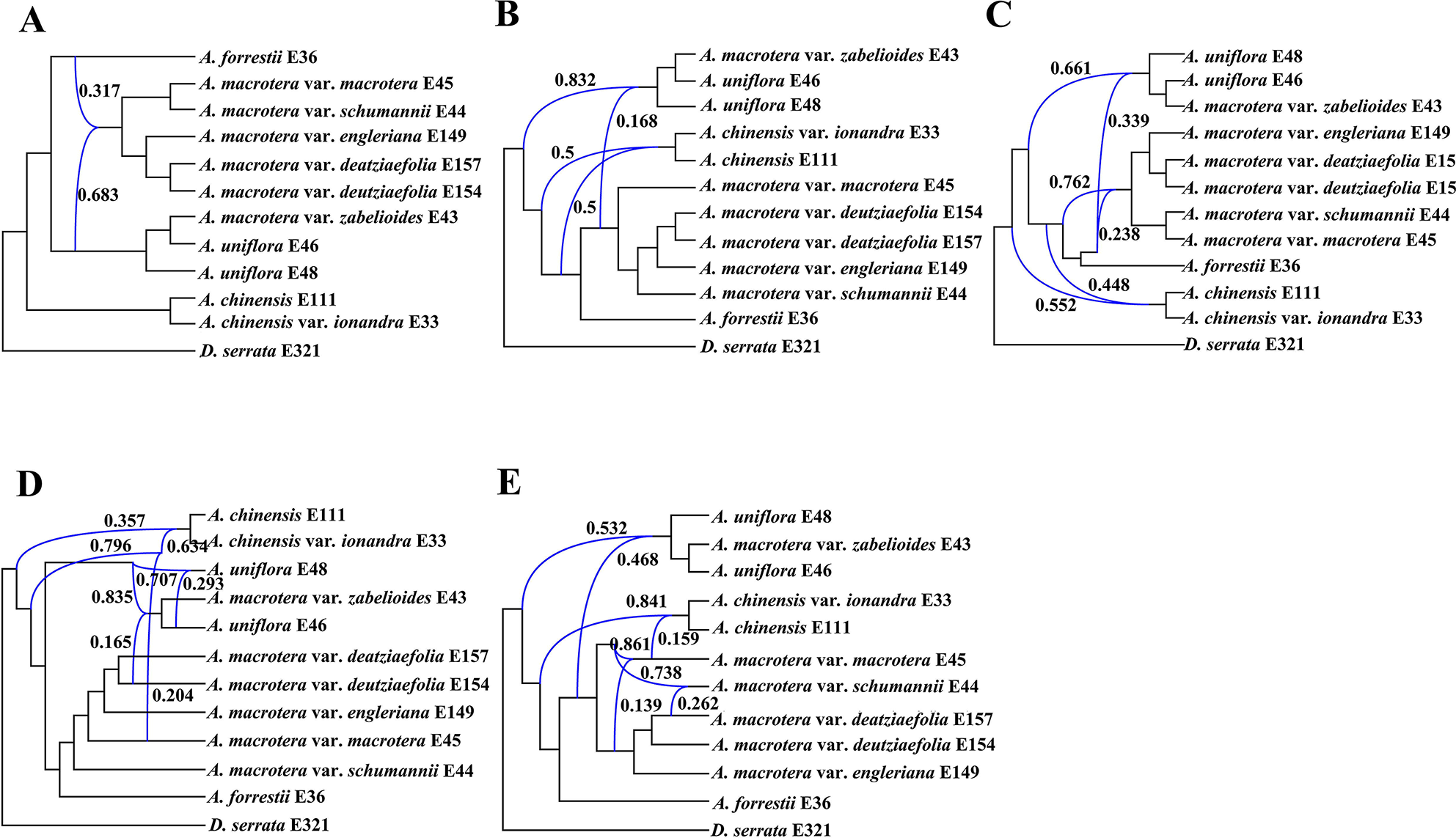
Best species networks of the selective nuclear data estimated with PhyloNet for the 12 samples data set. A: One hybridization event; B: Two hybridization events; C: Three hybridization events; D: Four hybridization events; E: Five hybridization events. Blue branches connect the hybrid nodes. Numbers next to blue branches indicate inheritance probabilities.

**Fig. S6.**
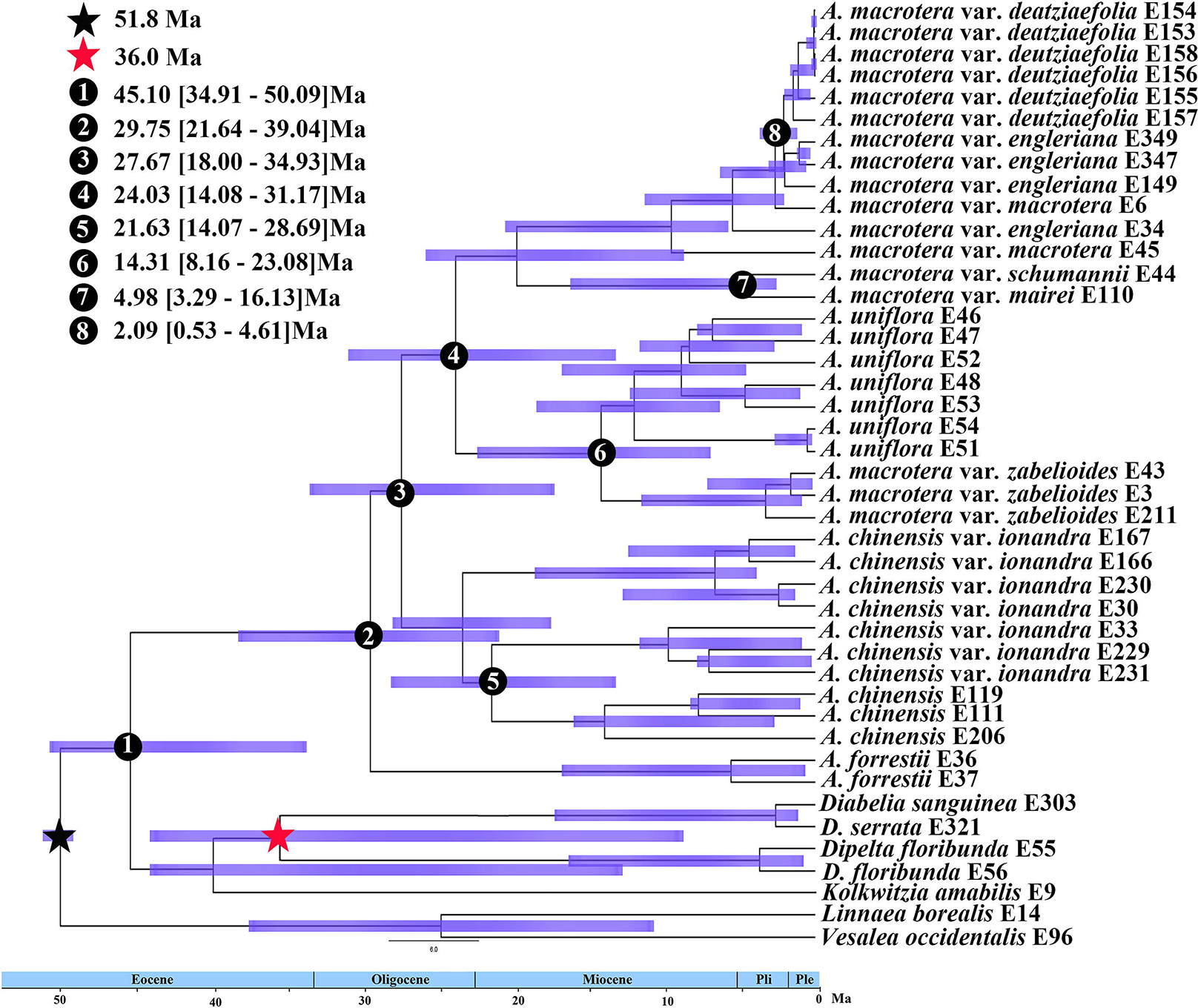
BEAST analysis of divergence times based on the plastid matrix. Calibration points are indicated by stars. Numbers 1-8 represent major divergence events in *Abelia*; Mean divergence times and 95% highest posterior densities are provided for each.

**Fig. S7.**
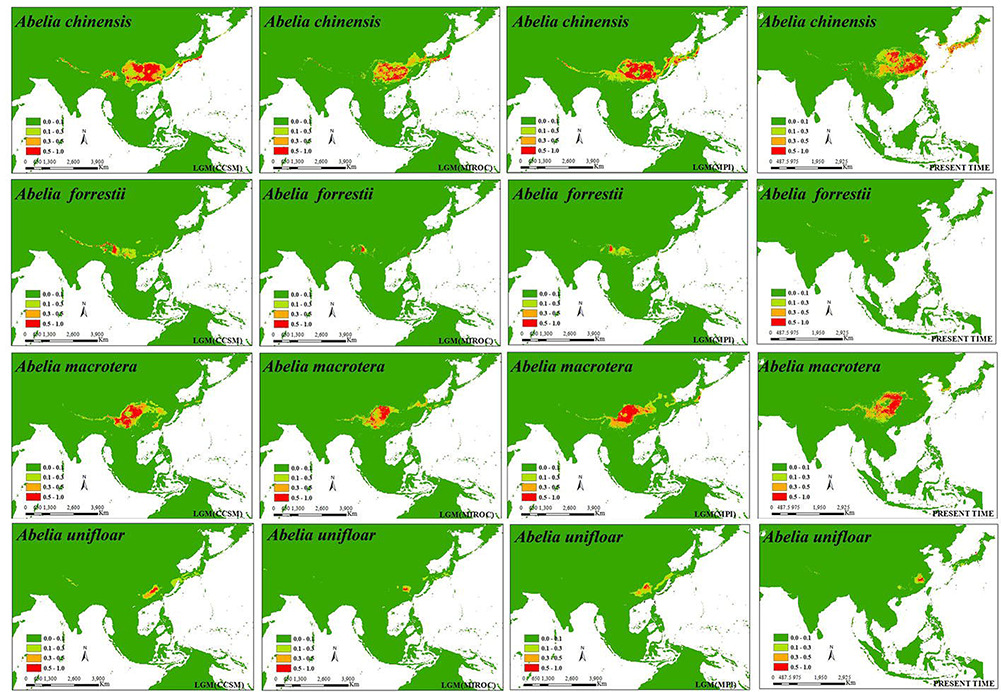
Potential distribution probability (in logistic value) of occurrence for *Abelia* in East Asia generated form MAXENT.

**Table. S1.**
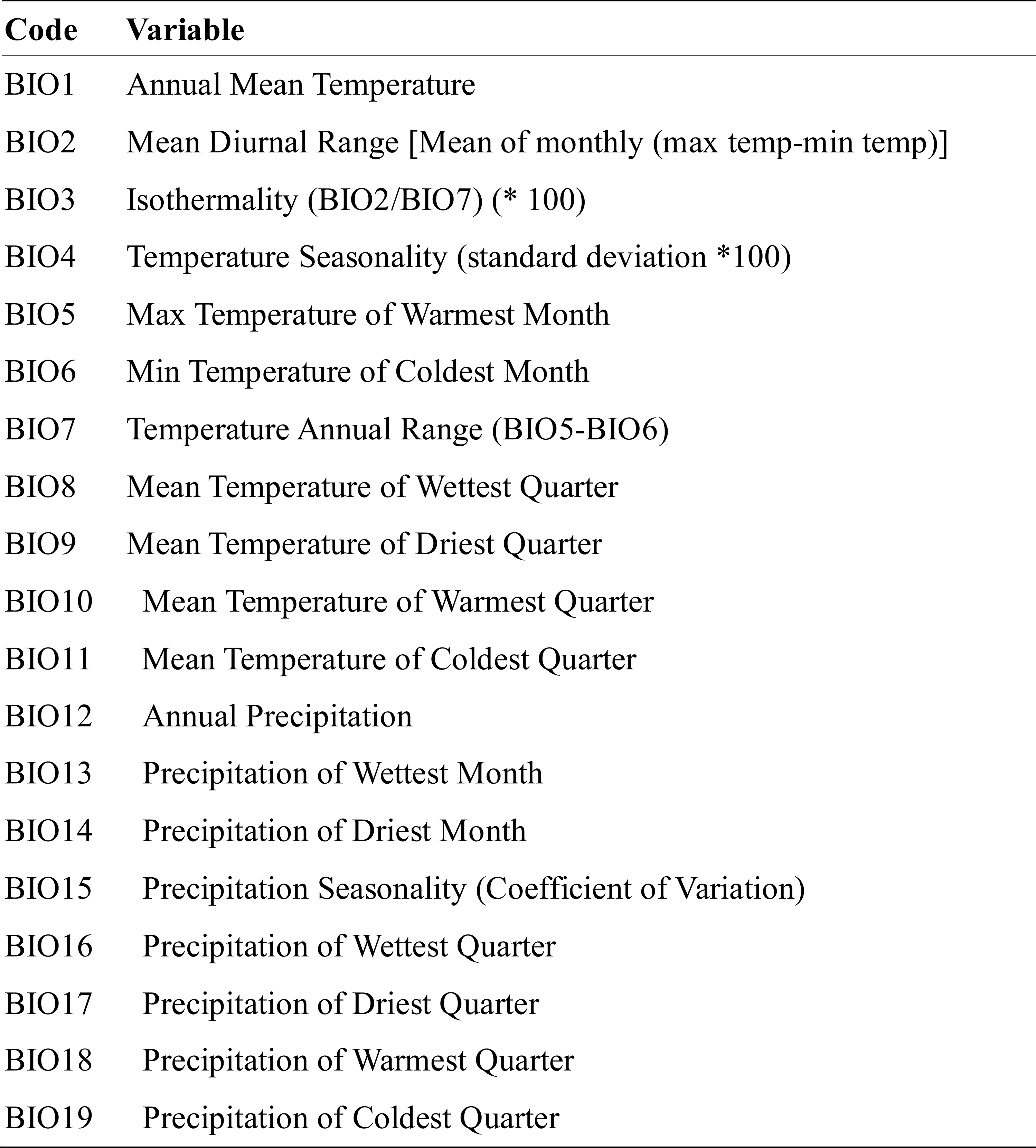
The 19 bioclimatic variables for ENM analyses

**Table S2.**
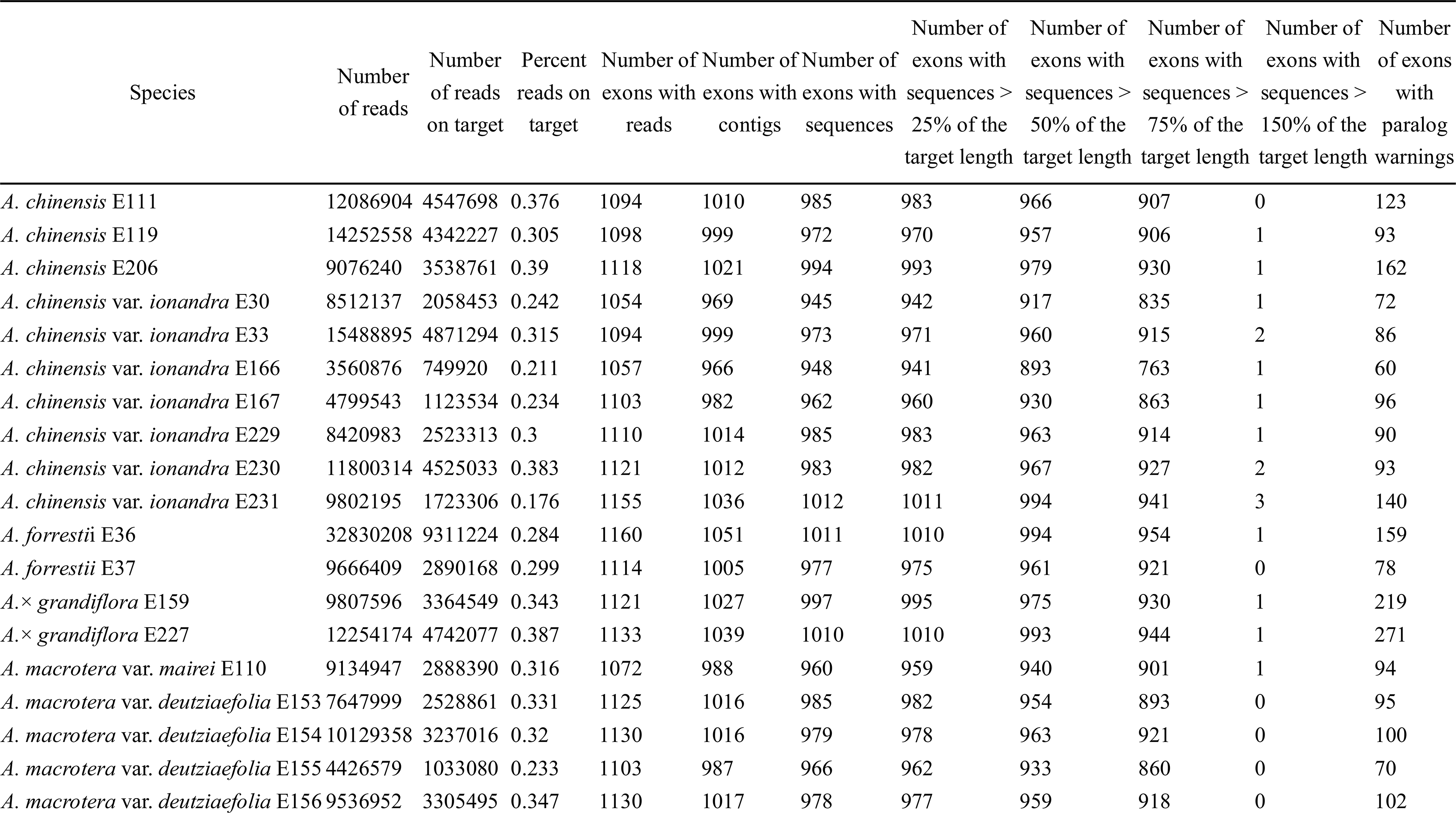
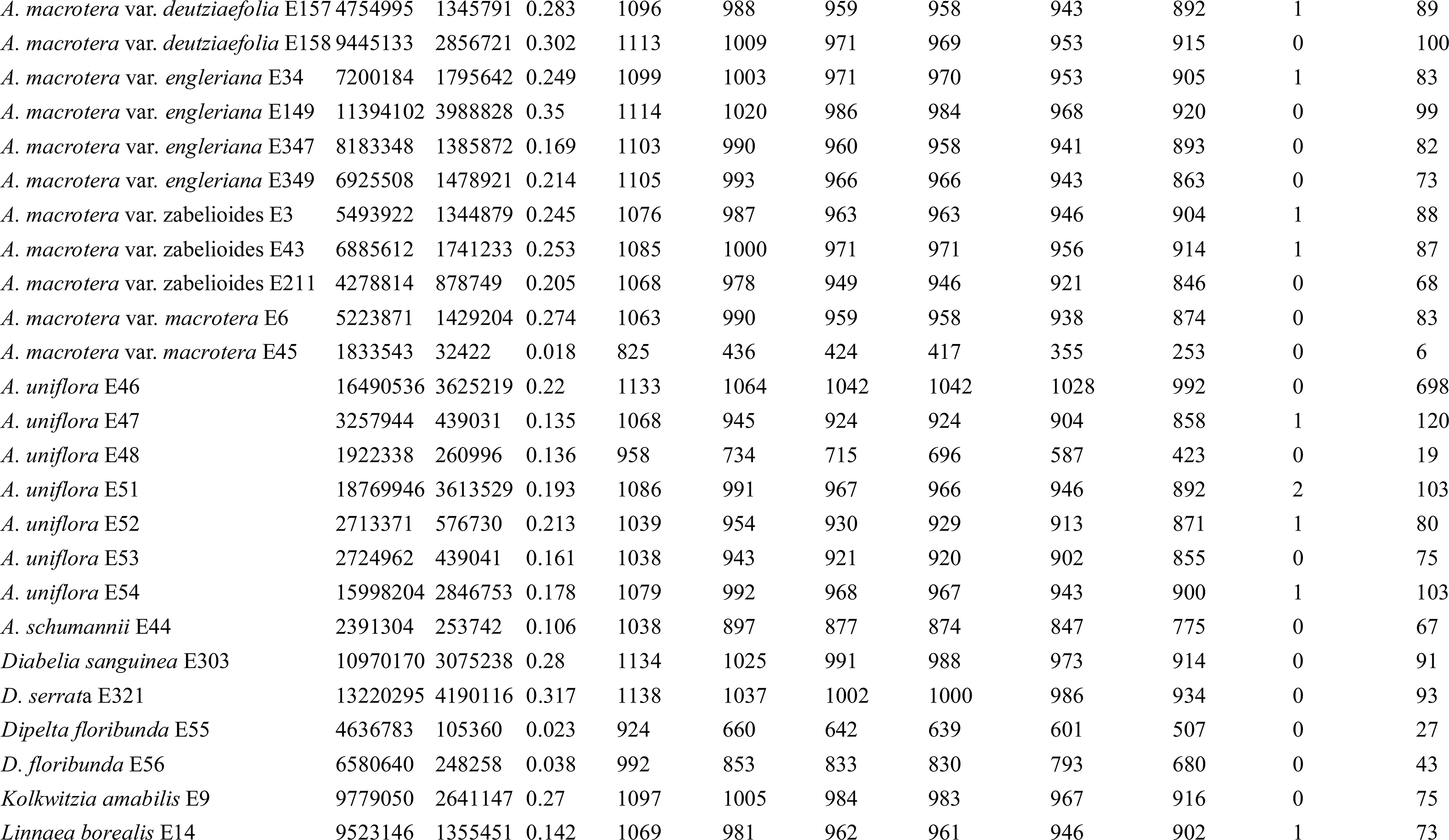
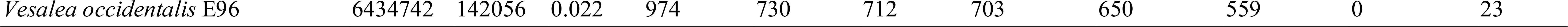
HybPiper assembly statistics

